# Ubiquitin ligase RNF126 exerts functions in lysosomal damage response and lysosomal homeostasis maintenance

**DOI:** 10.64898/2026.10.16.759566

**Authors:** Sophie Levantovsky, Bojana Kravić, Lizhen Cheng, Lukas Brecht, Hanna Tolle, Pinki Gahlot, Georg Kislinger, Kimberly Cramer, Regina Feederle, Benedikt Wefers, Wolfgang Wurst, Ralf Jungmann, Martina Schifferer, Hemmo Meyer, Christian Behrends

## Abstract

The response to lysosomal membrane permeabilization involves extensive ubiquitylation at the lysosome; however, the ubiquitin ligases identified so far cannot account for the complexity of the response. Using extensive spatial proteomics, we identified RNF126 as a new ligase that is robustly recruited to lysosomes in response to a range of different damage triggers across distinct cell types. Recruitment is mediated independent of BAG6 via an arginine-rich stretch in the low-complexity region of RNF126 to mostly unacidified lysosomes in the perinuclear region. Ubiquitin remnant mass spectrometry identified several damage-induced RNF126 substrates, among them known lysosomal damage response components and potential new factors that might drive regeneration from lysosomal damage. Consistently, RNF126 deletion resulted in persistent lysosomal swelling. In addition, we uncovered a constitutive role of RNF126 in lysosomal homeostasis maintenance. Hence, our findings identified the ubiquitin ligase RNF126 as novel lysosomal health regulating factor with functions in homeostasis and stress conditions.

## INTRODUCTION

Lysosomes represent the terminal compartment for numerous degradative and recycling pathways. Lysosomal membrane permeabilization (LMP) resulting in functional impairment, loss of acidic pH and leakage of lysosomal content can therefore critically impair cellular homeostasis and eventually lead to cell death.^1–3^ LMP can be induced by various triggers, including exposure to membranolytic agents, lipid peroxidation and unbalanced lysosomal lipid composition as well as uptake of proteotoxic aggregates or intracellular pathogens.^2, 4, 5^ Loss of lysosomal integrity is a feature in many pathophysiological conditions including atherosclerosis, neurodegeneration and lysosomal storage diseases.^4–6^ Understanding the cellular response to lysosomal injury may thus provide a strategy for identifying novel therapeutic targets.

Rapid responses to lysosomal membrane rupture involve ESCRT III-mediated membrane resealing as well as lipid replenishment via phosphoinositide-mediated ER-lysosome contact sites.^7–12^ In parallel, damage-induced mTOR inactivation results in TFEB translocation to the nucleus, where it induces lysosomal gene expression boosting lysosomal biogenesis.^13,14^ Eventually, incurable lysosomes are degraded via a selective form of macroautophagy, termed lysophagy.^15–17^ Extensive ubiquitylation observed upon LMP is mainly considered to be associated with lysophagy initiation involving recruitment of ubiquitin (Ub)-binding autophagy receptors which promote phagophore formation around the damaged lysosome.^1, 18–20^ Our previous work, however, revealed many LMP-induced ubiquitylation targets to be non-lysosomal raising the question to what extent Ub exerts regulatory functions within the lysosomal damage response beside serving as an lysophagy eat-me-signal.^21^ Consistent with that, recent studies identified additional regeneration processes differing from degrading the lysosome as a whole including ATG8ylation of the lysosomal membrane, LRRK2-mediated lysosomal tubulation, TBC1D15-mediated autophagic lysosome reformation (ALR) and partial internalization of lysosomal membrane patches and lysosomal membrane proteins via microautophagy.^5, 22–26^

Several Ub E3 ligases have recently been identified in the context of lysophagy initiation including CUL4A-DDB1-WDFY1 and SCF^FBXO^^27^. Both ligase complexes were described to ubiquitylate LAMP2 with K48-linked chains driving lysophagy.^19, 20^ Further, the HECT E3 ligase ITCH was shown to triage damaged lysosomes to lysophagy via catalyzing K63-linked Ub chains on lysosomes.^27^

However, the E3 ligases identified so far fail to explain the complexity of LMP-induced ubiquitylation of lysosomal proteins and additional regulators. Therefore, we employed unbiased lysosomal proximity labeling as well as immuno-isolation of lysosomes coupled to quantitative mass spectrometry (MS) to uncover novel Ub E3 ligases involved in the lysosomal damage response. Using this dual spatial proteomics approach, we identified the RING-type E3 ligase RNF126 associated with damaged lysosomes and validated its lysosomal translocation in different cell types upon various lysosomal damage modalities. Furthermore, we showed that recruitment of RNF126 is mediated by an arginine-rich stretch in its intrinsically disordered region. RNF126 deficiency results in a robust decrease of damage-induced ubiquitylation at the lysosome, a predisposition to lysosomal damage and an impaired lysosomal regeneration without affecting lysophagy. Next to its response to lysosomal damage, we revealed that RNF126 contributes to maintenance of lysosomal homeostasis by regulating lysosomal abundance of solute carrier (SLC) transporters potentially through Ub-dependent sorting as well as mTOR complex association with lysosomes. Overall, this study uncovered previously undescribed roles for RNF126 in contributing to lysosomal health and protecting cells from consequences of LMP.

## RESULTS

### Spatial proteomics reveals recruitment of E3 ligase RNF126 to damaged lysosomes

To identify E3 ligases taking part in the endolysosomal damage response, we took an unbiased APEX2-based proximity biotinylation approach.^28^ Thereto, we generated HeLa cells stably expressing the lysosomal transmembrane protein TMEM192 C-terminally fused to the engineered peroxidase APEX2 and a Flag epitope tag (Figure 1A) and confirmed full length and correct localization by immunoblotting and immunofluorescence microscopy. (Figure S1A). To induce biotinylation of proteins in close vicinity to lysosomes, TMEM192-APEX2-expressing cells were incubated with biotin-phenol (BP) for 30 min and subsequently pulsed with H_2_O_2_ for 1 min. Following this treatment, we observed robust biotinylation at LAMP1-positive vesicles, defined as lysosomes (Figure S1B). Next, we confirmed biotinylation of lysosomal membrane proteins in BP-treated but not mock-treated TMEM192-APEX2 expressing cells via streptavidin pulldown followed by on-bead tryptic digestion and MS analysis (Figure S1C). Gene Ontology (GO) enrichment analysis of significantly enriched proteins revealed terms such as endosomal and lysosomal membranes reaffirmed strong specificity towards lysosomal proximity (Figure S1D). Interestingly, we also detected an enrichment for ER-resident and cytosolic proteins reflecting the capability to capture transient proximity partners next to lysosomal membrane proteins.

**Figure 1.**
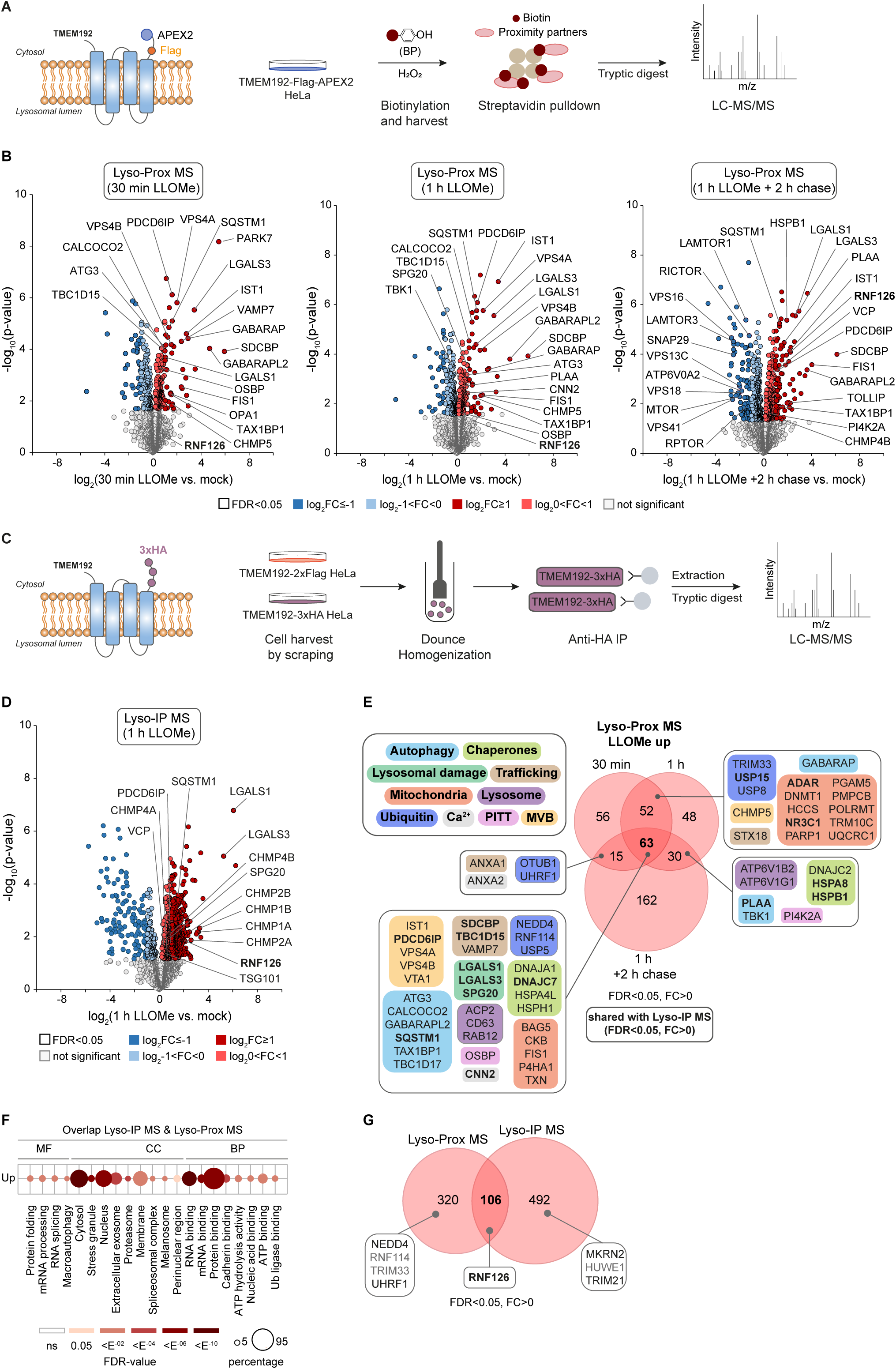
Spatial proteomics reveals recruitment of E3 ligase RNF126 to damaged lysosomes (A) Proximity biotinylation of lysosomes and quantitative mass spectrometry (Lyso-Prox MS) upon LLOMe treatment using TMEM192-Flag-APEX2 cells. (B) Volcano plots of lysosomal proximity proteome changes upon damage. Cells were treated with LLOMe over a time course of 30 min, 1 h and 1 h followed by a 2 h recovery phase (chase). Experiments were performed in biological quadruplicates and statistical analysis using two-sided two sample Student’s t-test (p-value<0.05) with permutations-based correction (FDR<0.05). Significantly altered proteins (FDR<0.05) are labeled in dark (log_2_FC ≥ I1I) and light color (log_2_I0<FC<1I), respectively. (C) Proteomics workflow for Lyso-IP MS using TMEM192-3xHA cells. (D) Volcano plot of lysosomal proteome changes. Cells were treated with LLOMe for 1 h followed by Lyso-IP MS. Experiments were performed in biological quadruplicates and statistical analysis using two-sided two sample Student’s t-test (p-value<0.05) with permutations-based correction (FDR<0.05). Significantly altered proteins (FDR<0.05) are labeled in dark (log_2_FC ≥ I1I) and light color (log_2_I0<FC<1I), respectively. (E) Manual annotation of common proteins significantly enriched in the proximity of damaged lysosomes (FDR<0.05, log_2_FC>0) at indicated time points from (B). Proteins also found significantly enriched (FDR<0.05, log_2_FC>0) using Lyso-IP from (D) are marked in bold. (F) GO enrichment analysis of proteins significantly enriched at lysosomes (FDR<0.05, log_2_FC>0) identified by Lyso-Prox (B) and Lyso-IP (D). Dot size correlates to number of proteins, dot color to term enrichment (FDR). BP, biological process; CC, cellular component, MF, molecular function. (G) Overlap of proteins significantly enriched at damaged lysosomes (FDR<0.05, log_2_FC>0) identified by Lyso-Prox (B) and Lyso-IP (D). Ub E3 ligases identified in both approaches are highlighted black. Grey shows Ub E3 ligases only detected in one of the two MS approaches. Ub E3 ligases found significantly enriched in both approaches are marked in bold.

To introduce lysosomal damage, we treated cells with the well-characterized lysosomotropic, membrane-permeable small molecule L-leucyl-L-leucine methyl ester (LLOMe). The mode of action of this dipeptide is a cathepsin C catalyzed conversion to a membranolytic polymer that induces robust permeabilization of late endosomal and lysosomal limiting membranes.^15, 29, 15, 30^ Our TMEM192-APEX2-expressing cells responded robustly to LLOMe-induced lysosomal damage at a modest concentration of 250 µM visualized by the appearance of Gal3 (alias LGALS3) on biotin-labeled lysosomes (Figure S1B). To monitor proteins recruited to lysosomes at early and advanced stages of lysosomal damage, we compared untreated TMEM192-APEX2 cells to those treated with LLOMe for 30 min, 1 h or 1 h followed by a 2 h recovery phase (chase), respectively. By coupling proximity labeling with quantitative mass spectrometry, we identified multiple protein classes recruited significantly to ruptured lysosomes at one or more time points post LLOMe, including well-characterized lysosomal damage markers (LGALS1, LGALS3, SPG20), ATG8 proteins (GABARAP, GABARAPL2), Ub-binding autophagy receptors (CALCOCO2, SQSTM1, TAX1BP1, TOLLIP) and proteins shown previously to be crucially involved in lysophagy (PLAA, VCP, TBK1, SPG20) (Figures 1B, 1E, Table S1).^27, 31, 32, 27^ In concert with previous studies, we also detected a strong enrichment of components involved in ESCRT-mediated repair (CHMP5, CHMP4B, IST1, VPS4B, PDCD6IP, VPS4A) as well as components of the phosphoinositide-initiated membrane tethering and lipid transport (PITT) pathway (PI4K2A, OSBP) for rapid lysosomal repair.^7–9, 11, 12^ Consistently, we find ESCRT-driven pathways, e.g. multivesicular body (MVB) assembly and endocytosis, strongly enriched in GO term analysis next to autophagy upon acute damage (Figures S1E, S1F). Of note, a robust dissociation of mTORC1 (MTOR, RPTOR, LAMTOR1, LAMTOR3, RICTOR) and HOPS (VPS16, VPS18, VPS41) complexes from lysosomes as well as disassembly of the v-ATPase was observed during the recovery phase (Figures 1B, S1G, Table S1). The loss of mTORC1 kinase complex from the lysosomal surface was previously described to be Gal8-mediated, however, we were unable to capture LGALS8 at damaged lysosomes in our cellular model.^13^ Among the Ub machinery, we identified several DUBs (OTUB1, USP5, USP8, USP15), one E2 conjugating enzyme (UBE2O) and five E3 ligases (NEDD4, RNF114, TRIM33, UHRF1, RNF126) in at least one of the time points post LLOMe (Figures 1B, 1E, Table S1). Proteins of the TRIM (tripartite motif) family were shown to interact with Gal3 and for TRIM16 in particular a role as recruitment factor for autophagy regulators ULK1 and Beclin 1 in response to lysosomal damage was reported.^33^ The HECT ligase NEDD4 was described to regulate sorting and degradation of short-lived lysosomal membrane proteins via ubiquitylation followed by ESCRT-mediated intraluminal invagination.^34^ Notably, the three RING ligases RNF114, UHRF1 and RNF126 have not been linked to damaged lysosomes in any way.

To validate these results, we complemented our proximity labeling approach with the more stringent Lyso-IP method using TMEM192-3xHA expressing HeLa cells.^35^ The specificity of lysosome purification was benchmarked with TMEM192-2xFlag expressing control cells. Expression of both fusion proteins was confirmed via immunoblotting (Figure S2A). For Lyso-IP, cells were dounce-homogenized, lysosomes isolated with anti-HA magnetic beads and eluted with denaturing Urea buffer. Lysosomal proteins were subsequently precipitated, digested with trypsin and analyzed by MS (Figure 1C, Table S2). Statistical analysis of HA-vs. Flag-tagged TMEM192 expressing cells showed prominent enrichment of lysosomal membrane proteins as well as luminal proteins in contrast to APEX2-driven proximity proteomics which only enables cytosolic labeling due to membrane-impermeability of BP (Figure S2B). In line with that, we observed prominent lysosome-associated GO annotations involving proteolytic and hydrolytic activities, confirming proper isolation of lysosomes and their content (Figure S2C). Comparing damaged and healthy lysosomes after immune-isolation revealed the accumulation of damage markers (LGALS1, LGALS3, SPG20), autophagy-related proteins (VCP, PLAA, SQSTM1) as well as ESCRT machinery components (PDCD6IP, CHMP1A, CHMP1B, CHMP4A, CHMP4B, TSG101) (Figure 1D, S2D, Table S2) in line with proximity proteomics and previous studies.^8, 32, 8^ With regard to Ub E3 ligases, we detected MKRN2, HUWE1, TRIM21 and RNF126 that were significantly enriched on damaged lysosomes. As expected, proteins detected by both spatial proteomics approaches shared GO terms such as perinuclear region, macroautophagy and Ub ligase binding (Figures 1F). Among this pool of 106 proteins, RNF126 was the only damage-responsive E3 ligase found significantly enriched at lysosomes (Figure 1G). Other E3 ligases were either only identified by one of the two approaches (RNF114, TRIM33, HUWE1) or were not significantly altered (NEDD4, UHRF1, MKRN2, TRIM21). In summary, APEX2-based proximity biotinylation and Lyso-IP both led to robust identification of known and new lysosomal damage factors amongst them RNF126.

### RNF126 translocates to lysosomes in a range of damage settings and cell types

To validate RNF126 as new component of the lysosomal damage response, we generated HeLa cells stably expressing N-terminally HA-tagged RNF126. Consistent with our proteomics results, LLOMe treatment induced pronounced translocation of cytosolic RNF126 to lysosomes (Figure 2A). Notably, neither the expression level of the fusion construct nor the abundance of the lysosomal transmembrane protein LAMP1 were affected by the treatment (Figure S3A). Nevertheless, LAMP1-positive structures became enlarged in line with previous reports.^36^ To determine the distribution of RNF126 at damaged lysosomal membranes in more detail, we monitored GFP-RNF126 expressing cells using DNA-PAINT technology and super-resolution microscopy (Figure 2B). Upon LLOMe treatment, lysosomes showed a clear dysmorphology, lost their round shape and occupied a bigger area. While RNF126’s extensive colocalization with lysosomes was confirmed under this condition, RNF126 was not found to be uniformly distributed but instead showed a nanoscale patch-like localization pattern on damaged lysosomes, possibly indicating recruitment of RNF126 to distinct subdomains of the lysosomal membrane.

**Figure 2.**
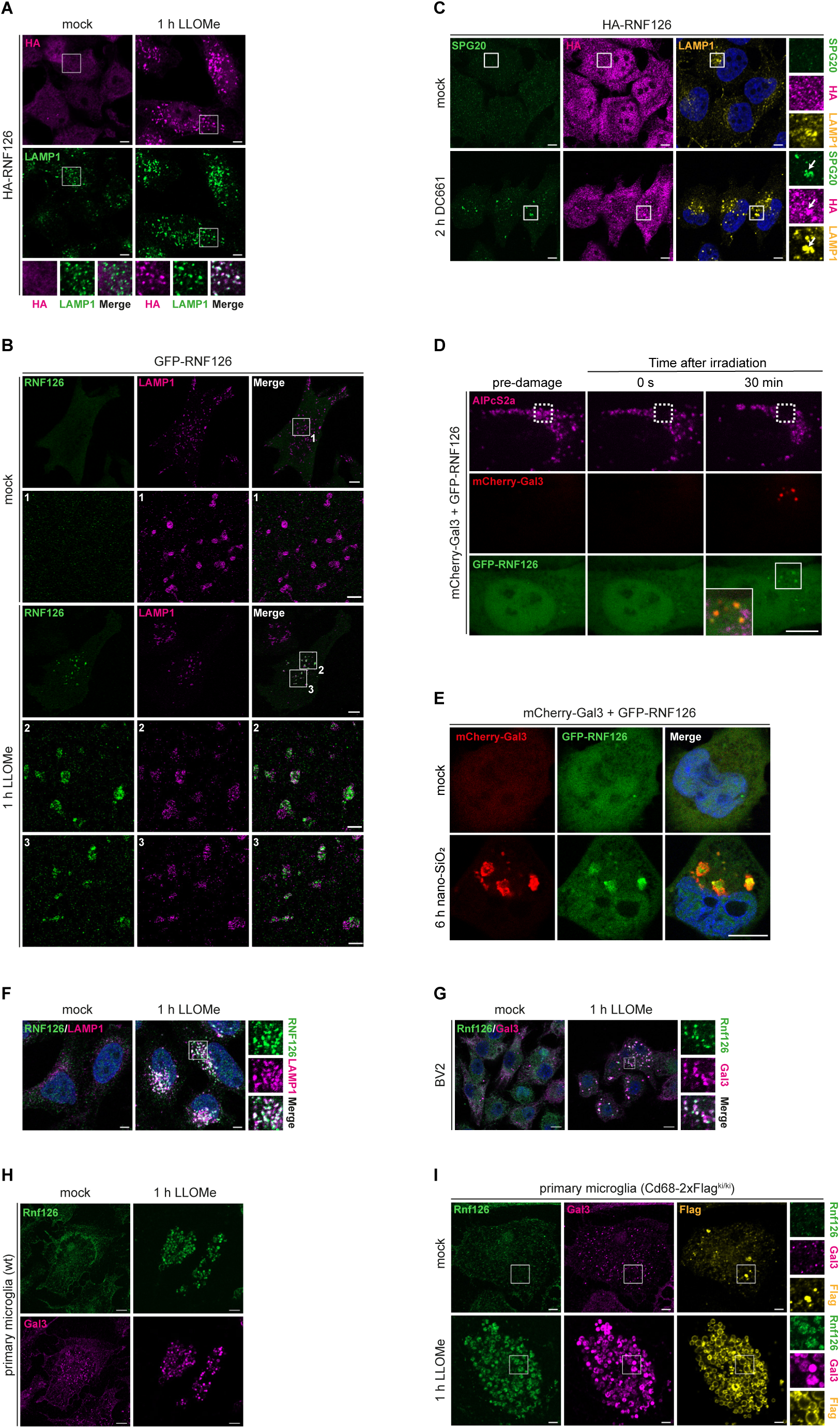
Lysosomal translocation of RNF126 is conserved across different damage and cell types (A) Cells expressing HA-RNF126 were treated with LLOMe, fixed, stained with anti-HA and -LAMP1 antibodies and subjected to confocal microscopy. Insets show magnification of boxed areas. Scale bars, 5 µm. (B) Cells expressing GFP-RNF126 were treated with LLOMe, fixed, immunolabeled with an anti-LAMP1 antibody and subjected to DNA-PAINT technology for super-resolution microscopy. Scale bars, 5 µm. Insets show magnification of boxed areas. Scale bars of insets, 1 µm. (C) Cells expressing HA-RNF126 were treated with the PPT1 inhibitor DC661 (2 h, 3 µM), fixed and stained with indicated antibodies. Insets show magnifications of boxed areas. Arrow indicates colocalization. Scale bars, 5 µm. (D) GFP-RNF126 and mCherry-Gal3 expressing cells were loaded with photosensitizer AlPcS2a, irradiated in the indicated area and chased over the course of 30 min by live cell microscopy. Scale bars, 5 µm. (E) GFP-RNF126 and mCherry-Gal3 expressing cells were either mock treated or incubated with nano-silica (50 µg/ml) for 6 h before fixation. Scale bars, 5 µm. (F) Cells were treated with LLOMe, fixed and stained with anti-RNF126 and -LAMP1 antibodies. Insets show magnification of boxed areas. Scale bars, 5 µm. (G) Murine microglial-like cells (BV2) were treated with LLOMe (2 mM), fixed and stained with anti-Rnf126 and -Gal3 antibodies. Insets show magnification of boxed areas. Scale bars, 10 µm. (H) Primary microglia isolated from 8-months-old wild-type mice and cultured for 7 days were treated with LLOMe (500 µM), fixed and stained with anti-Rnf126 and -Gal3 antibodies. Scale bars, 10 µm. (I) Primary microglia isolated from 8-months-old Cd68-Flag^KI/KI^ mice and cultured for 7 days were treated with LLOMe (500 µM), fixed and stained with indicated antibodies. Insets show magnification of boxed areas. Scale bars, 10 µm.

Importantly, treatment with the lysosomotropic damaging agent glycyl-l-phenylalanine 2-naphthylamide (GPN), known to induce osmotic stress and subsequent lysosomal membrane rupture^37^, led to a translocation of RNF126 to Gal3-positive lysosomes (Figure S3B). In contrast, milder lysosomal perturbation, such as inhibition of lysosomal V-ATPase with Bafilomycin A1 or treatment with conjugation of ATG8 to single membrane (CASM) inducer monensin^38–40^ were insufficient to recruit RNF126 to lysosomes (Figures S3C, S3D). We therefore tested harsher perturbations, such as induction of lysosomal lipid peroxidation (LLP) that results in lipid-packaging defects due to lipid truncation and the extra space taken by lipid hydroperoxides.^41, 42^ We induced lipid peroxidation in several ways. Firstly, we treated HA-RNF126 expressing cells with the palmitoyl-protein thioesterase 1 (PTT1) inhibitor DC661 which triggered recruitment of HA-RNF126 to lysosomes marked with lipid-packaging defect marker SPG20 (Figure 2C).^27, 43^. Secondly, we generated radicals by irradiation of photosensitizer AlPcS2a–loaded lysosomes (Figure 2D)^16, 21^ or by uptake of nano-silica beads.^44^ Importantly, both treatments triggered prominent recruitment of GFP-RNF126 to mcherry-Gal3-decorated, damaged lysosomes (Figures 2D, 2E).

Next, we tested for lysosomal recruitment of endogenous RNF126. Using custom-made antibodies, we found human and murine RNF126/Rnf126 strongly recruiting to damaged lysosomes in HeLa and murine BV2 microglial like cells, respectively (Figures 2F, 2G). We also checked for Rnf126 recruitment in primary microglia isolated from adult mice brains. These highly specialized, phagocytic cells were challenged with LLOMe after one week in culture. In these cells, we detected striking translocation of Rnf126 to Gal3-positive damaged lysosomes (Figures 2H). We further made use of a Cd68-Flag^ki/ki^ knock-in mouse model, where microglial-specific lysosomal transmembrane protein Cd68 is fused to a triple Flag tag, and observed colocalization of Cd68, Gal3 and Rnf126 upon LLOMe treatment (Figure 2I). Together, these results show that the Ub E3 ligase RNF126 translocates to lysosomes upon various damage triggers ranging from membranolytic lysosomotropics to lipid peroxidation and that this translocation is conserved across diverse human and mouse cell models.

### RNF126 is recruited to lysosomes after initial damage recognition

To start dissecting RNF126’s function within the lysosomal damage response, we analyzed the recruitment dynamics of RNF126 in comparison to several established components of this pathway. Briefly, the ESCRT repair machinery component IST1 showed fast recruitment which was followed with a delay by SPG20, Gal3 and LC3 in line with previous reports (Figure 3A).^7, 27^ Notably, the majority of ATG8-lipidation observed upon damage has recently been shown to be due to CASM and distinct from initiation of lysophagy.^22^ Nevertheless, we saw an early translocation of the selective autophagy receptor p62 (also known as SQSTM1). However, its contribution to lysophagy or CASM remains poorly understood ^31, 32^ Intriguingly, RNF126 appears rather late after initial damage detection via ESCRT and translocation of the other damage response components, starting around 30 min post-LLOMe addition. This suggests that RNF126 unlikely contributes to ESCRT-mediated repair. Recently, an additional ESCRT-independent repair mechanism involving lipid-transfer from ER to lysosome and PI4K2A-catalyzed lipid conversion to phosphatidylinositol-4-phosphate (PI4P) was described.^11, 12^ We tested whether RNF126 is an effector of this pathway. However, PI4K2A KO cells showed no effect on RNF126 recruitment upon damage (Figures S3E and S3F), indicating that RNF126’s response to damaged lysosomes is uncoupled from the appearance of PI4P.

**Figure 3.**
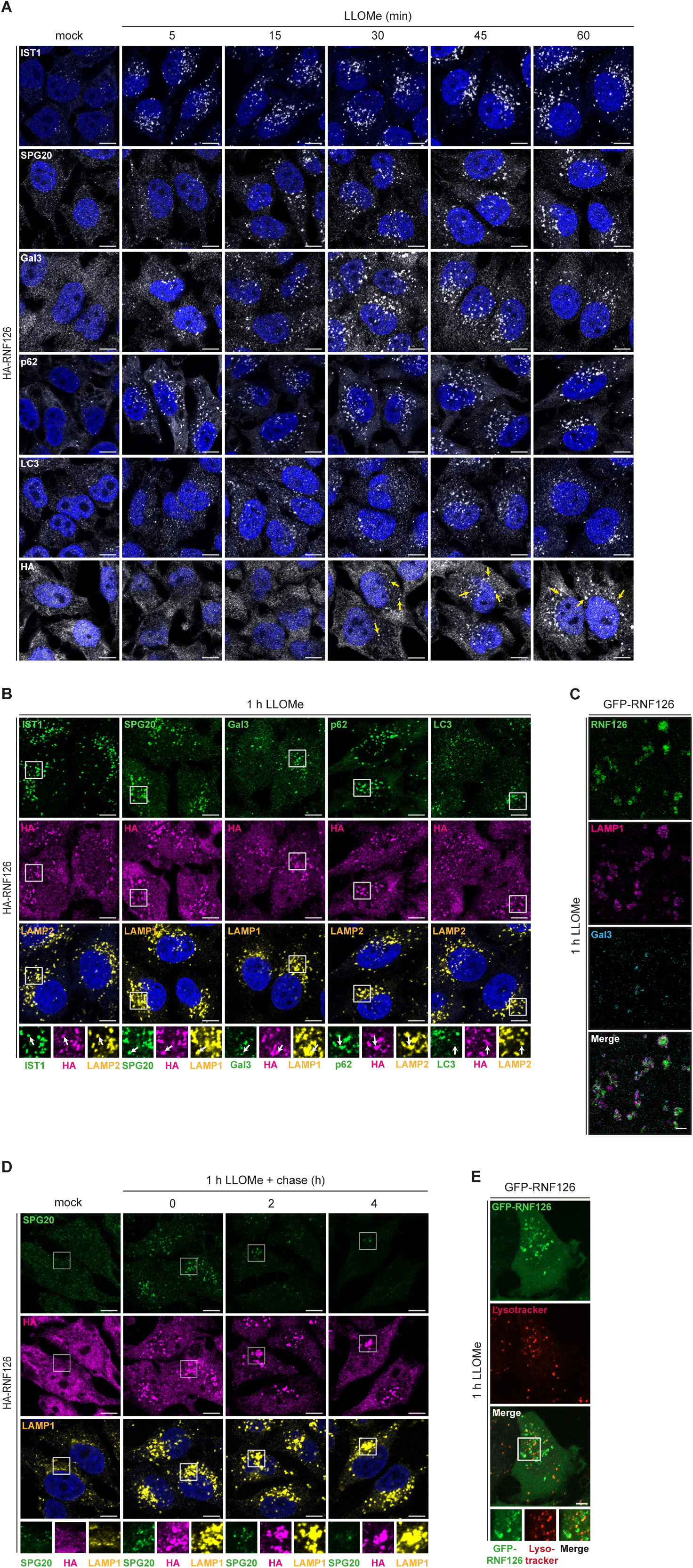
RNF126 is recruited to lysosomes after initial damage recognition (A) HA-RNF126 cells were treated up to 60 min with LLOMe, fixed and stained with indicated antibodies. Yellow arrows indicate RNF126 translocation to lysosomes. Scale bars, 10 µm. (B) HA-RNF126 cells were treated with LLOMe, fixed and stained with indicated antibodies. Insets show magnifications of boxed areas and arrows colocalization events. Scale bars, 10 µm. (C) GFP-RNF126 cells were treated with LLOMe, fixed, immunolabeled with anti-LAMP1 and-Gal3 antibodies and subjected to DNA-PAINT technology for super-resolution microscopy. Scale bars, 1 µm. (D) HA-RNF126 cells were treated with LLOMe followed by washout and 4 h chase with fresh media. Cells were fixed and stained with indicated antibodies. Insets show magnifications of boxed areas. Scale bars, 10 µm. (E) GFP-RNF126 cells were treated with LLOMe, stained with Lysotracker Red DND-99 and fixed. Scale bars, 10 µm.

Costaining of RNF126 with the above-mentioned markers revealed that RNF126 shows strong colocalization with IST1, SPG20 and p62, whereas LC3 and Gal3 decorated lysosomes only share partial overlap (Figure 3B). DNA-PAINT-based super-resolution microscopy confirmed that RNF126 and Gal3 label different lysosomal populations (Figures 3C, S3G), indicating different recruitment cues for these two damage response factors. To exclude a potential upstream role of Gal3, as described for the E3 ligase TRIM16, we monitored RNF126 translocation in Gal3 KO cells and found no dependency on Gal3 (Figures S3H, S3I).^33^ Similarly, p62 depletion did not affect RNF126 localization to lysosomes (Figures S3J, S3K).

Given RNF126’s late arrival at damaged lysosome, we thought to determine its lysosomal residence time. Thereto, we performed a pulse-chase experiment where HA-RNF126 expressing cells were treated with LLOMe for 1 h followed by up to 4 h recovery in fresh media. Interestingly, RNF126 persisted on perinuclear located lysosomes up to 4 h in contrast to SPG20 which dissociated completely within this time frame (Figure 3D). We next asked how RNF126-decorated lysosomes differed from the RNF126-negative population. To address this, we performed lysotracker fluorescence recovery experiments widely used in the field to distinguish between compromised and repaired lysosomes.^8, 27^ GFP-RNF126 expressing cells were loaded with lysosomotropic pH-sensitive Lysotracker Red DND-99 and treated with LLOMe for 1 h. We observed recovery of lysotracker signal, reflecting repaired and re-acidified lysosomes, but interestingly, RNF126 preferentially labeled lysotracker-negative lysosomes that failed to recover (Figure 3E). This result is in line with the long persistence of RNF126 on lysosomes and the robust co-appearance with SPG20 which was shown to label lysosomes that fail to recover swiftly from damage.^27^

### Lysosomal recruitment is mediated by the low complexity region of RNF126

To determine how RNF126 recognizes compromised lysosomes, we used AlphaFold for structure prediction. RNF126 comprises two functional domains: A C4-type Zinc finger motif at its N-terminus (aa 13-32) and the catalytical RING finger domain (aa 229-270) at its C-terminus (Figure 4A). The middle part of the protein is predicted to be of low complexity and disordered. To test whether RNF126’s ligase activity is required for recruitment to damaged lysosomes, we reconstituted CRISPR/Cas9-generated RNF126 KO cells (Figure 4B) with RNF126 wild-type or catalytic dead mutant (C229/232A).^45, 46^ However, the enzymatic activity of RNF126 was dispensable for its translocation (Figure 4C).

**Figure 4.**
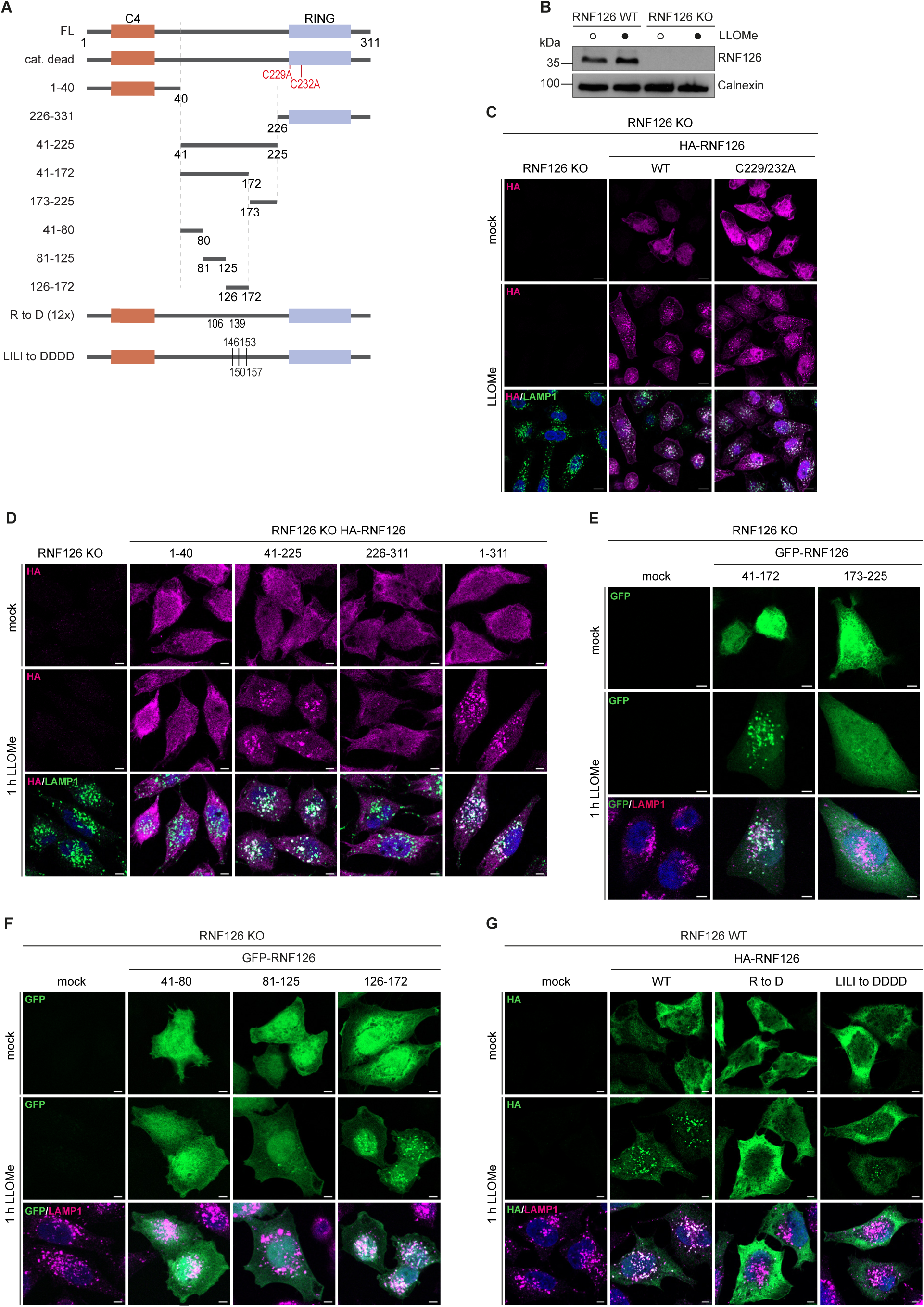
Lysosomal recruitment is mediated by the low complexity region of RNF126 (A) Scheme of RNF126 variants. (B) Immunoblot of lysates from RNF126 WT and KO cells. (C and D) RNF126 KO cells reconstituted with indicated HA-RNF126 variants were treated with LLOMe, fixed and stained with anti-HA and -LAMP1 antibodies. Scale bars, 10 µm (C) and 5 µm (D). (E and F) RNF126 KO cells expressing GFP-RNF126 variants were treated with LLOMe, fixed and stained with indicated antibodies. Scale bars, 5 µm. (G) Cells expressing indicated HA-RNF126 variants were treated with LLOMe, fixed and stained with anti-HA and -LAMP1 antibodies. Scale bars, 5 µm.

Previous reports showed that RNF126 is crucially involved in targeting cytosol-mislocalized membrane proteins with exposed hydrophobic stretches for proteasomal degradation.^47, 48^ Binding of these substrates was mediated by the chaperone BAG6 which in turn interacts with the N-terminal Zinc finger motif of RNF126.^47, 49^ We hypothesized that RNF126 exerts a similar function upon lysosomal damage in which BAG6 would bind defective lysosomal membrane proteins and recruit RNF126 to these clients to destine them for proteasomal degradation. To test this hypothesis, we designed truncations comprising of (i) the C4-type Zinc finger (aa 1-40), (ii) the disordered middle part of the protein (aa 41-225) and (iii) the catalytic RING domain (aa 226-311) (Figure 4A). We stably introduced these truncations fused to an HA tag into RNF126 KO cells and confirmed their similar expression via immunoblotting (Figures S4A). While the Zinc finger was sufficient to bind BAG6 (Figure S4B) as independently shown by a NMR study ^49^, this part of RNF126 was unable to translocate to lysosomes upon LLOMe (Figure 4D). Since BAG6 neither was found to be damage-enriched at lysosomes in our proximity or organelle proteomics data sets (Table S1 and S2) nor did BAG6 knockdown (KD) affect RNF126 recruitment to damaged lysosome (Figures S4C, S4D), we concluded that lysosomal translocation of RNF126 is not driven by BAG6.

Consistent with the finding that the catalytic activity is dispensable for lysosomal recruitment, the RING domain alone was unable to translocate to lysosomes (Figures 4C, 4D). Surprisingly, the middle part of RNF126 which does not comprise any high confidence structural features and was previously not examined in any other context was sufficient to trigger translocation upon LLOMe (Figure 4D). Splitting the middle part into two consecutive fragments (Figures 4A) revealed that fragment aa 41-172 is sufficient to drive translocation (Figures 4E). Analysis of this region by three additional fragments (Figures 4A) showed that aa 126-172 exhibited the strongest recruitment whereas aa 81-125 showed a markedly reduced but not completely abolished translocation (Figure 4F). Based on closer examination of the aa composition of these two stretches, we hypothesized that RNF126 might associate directly with the lysosomal membrane through either a poly-arginine stretch (aa 106-139, 12 arginine residues) which binds anionic lipids in the lysosomal lipid bilayer or an amphipathic helix (aa L146, I150, L153, L157; LILI motif) (Figure S4E). To test these hypotheses, we disrupted both motifs by aspartate point mutations to reverse the charge of arginine residues and to abolish the hydrophobicity for the LILI motif (Figure 4A). While mutation of LILI to DDDD strongly reduced but not abolished recruitment, disrupting the positive charge in the R to D mutant completely suppressed translocation (Figure 4G). Given that RNF126 translocation is induced by lysosomal lipid peroxidation (Figure 2C-E), we hypothesize that RNF126 may form electrostatic interactions with peroxidized lipid species via its arginine-rich region (aa 106–139). However, this interaction might further be stabilized by hydrophobic interactions involving the amphipathic helix of RNF126.

### RNF126 contributes to damaged-induced ubiquitylation signature

Next, we asked whether RNF126 contributes to ubiquitylation observed following lysosomal damage. As an initial approach, we examined the temporal dynamics of lysosomal ubiquitylation relative to RNF126 recruitment along with potential colocalization between Ub and RNF126. Thereto, we performed a time-course analysis monitoring the early response from 5 minutes up to 60 minutes after LLOMe treatment. K48-linked Ub chains began to appear around 30 minutes. Intriguingly, RNF126 not only exhibited a similar recruitment pattern but also colocalized with K48-linked Ub chain-positive lysosomes (Figure 5A). Given that RNF126 is capable of catalyzing both K48-and K63-linked Ub chains, we next monitored K63 Ub on lysosomes and observed a comparable pattern to that of K48 Ub (Figure 5B).^46^ Since RNF126 persists on damaged lysosomes over time, we extended our analysis to later stages of the lysosomal damage response using pulse-chase experiments as before. Similar to earlier time points, RNF126 strongly colocalized with both K48 and K63 Ub chains (Figures 5C, 5D). These findings demonstrate that RNF126 recruitment coincides temporally with the appearance of K63 and K48 Ub chains at damaged lysosomes, suggesting an active role for RNF126 in damage-induced ubiquitylation. Consistent with this notion, we observed a marked reduction in LLOMe-induced K48-linked Ub across all time points in RNF126 KO cells (Figures 5E, 5F). Importantly, overall ubiquitylation levels remained unchanged in the absence of RNF126 and were not affected by LLOMe treatment (Figures S4F, S4G). Together, these results show that RNF126 contributes to damage-induced ubiquitylation at lysosomes.

**Figure 5.**
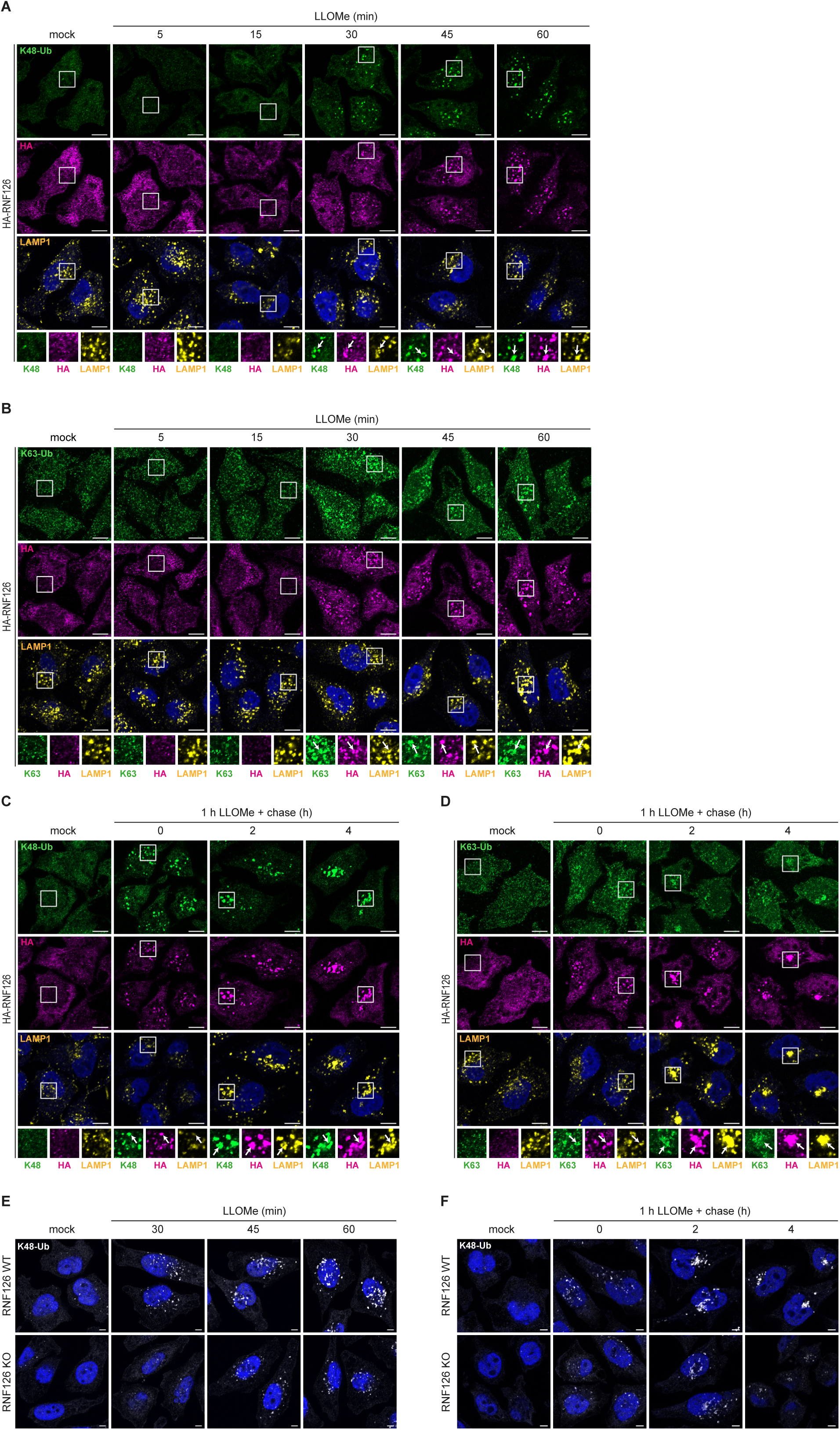
RNF126 contributes to damaged-induced ubiquitylation signature (A and B) HA-RNF126 cells were treated up to 60 min with LLOMe, fixed and stained with indicated antibodies. Insets show magnifications of boxed areas. White arrows indicate colocalization events. Scale bars, 10 µm. (C and D) HA-RNF126 cells were treated with LLOMe followed by washout and 4 h case with fresh media. Cells were fixed and stained with indicated antibodies. Insets show magnifications of boxed areas. White arrows indicate colocalization. Scale bars, 10 µm. (E) RNF126 WT and KO cells were treated up to 60 min with LLOMe, fixed and stained with a K48 Ub antibody. Scale bars, 5 µm.

### RNF126 substrates possess diverse functions associated with damaged lysosomes

To identify RNF126 substrates in response to lysosomal damage, we employed quantitative diGly proteomics. For this purpose, we combined stable isotope labeling by amino acids in cell culture (SILAC) with immunoaffinity-based enrichment of Ub remnant-containing peptides after tryptic digestion of ubiquitylated proteins. Using a two-step strategy, we first assessed LLOMe-induced ubiquitylation in parental cells (Figure S5A) and then compared RNF126-deficient and parental cells both treated with LLOMe (Figure S5B). Proteins that showed both a significant increase in ubiquitylation upon LLOMe treatment in parental cells and a significant decrease in LLOMe-treated RNF126 KO cells were considered potential RNF126 substrates. In the first step, we observed pronounced LLOMe-induced ubiquitylation which was not restricted to lysosomal proteins (TMEM192, LAMTOR1, PPT1) but also affected numerous non-lysosomal proteins involved in lysosomal damage-associated cellular processes including stress granule formation (DDX3X, PCBP2, UBAP2L, TARDBP (also known as TDP-43)), vesicular trafficking (CLINT1, PICALM, AGFG1), actin modulation (CNN2, CNN3), lipid metabolism (PLIN3, CAV1) and necroptotic death signaling (MLKL, RIPK1) (Figure 6A, Table S3). The second step of our analysis revealed marked impact of RNF126 deficiency on lysosomal damage-associated ubiquitylation as many of the aforementioned protein classes showed reduced ubiquitylation in LLOMe-treated RNF126 KO cells (Figure 6B, Table S4). Importantly, both datasets showed a striking overlap (Figure 6C) with several proteins (CNN2, PCBP2, DDX3X, ANXA2, TARDBP, PICALM, MLKL) harboring common ubiquitylation sites that were found regulated in a diametrically opposed manner dependent on LLOMe and RNF126 (Figures 6A, 6B) and that did not result from changes in protein abundance (Figure S5C). The broad effect of LLOMe-induced ubiquitylation on non-lysosomal proteins as well as the regulation of these events by RNF126 strongly supports the notion that ubiquitylation is not solely serving as degradative signal to drive lysophagy but exhibits additional regulatory functions. For instance, stress granule formation was recently discovered to be induced at damaged lysosomes and have a protective function by stabilizing the damaged lysosomal membrane, a function similar to ESCRTs that form filamentous spindles.^50, 51^ Another example is the RAB7 GTPase-activating protein TBC1D15 whose ubiquitylation was induced by damage and depended on RNF126. Recently, TBC1D15 was described as a scaffold protein for the assembly of autophagic lysosomal reformation machinery, a TFEB-independent regeneration pathway differing from lysophagy.^24^ An interesting neurodegenerative disease-linked protein that we identified as potential RNF126 substrate at damaged lysosomes is TDP-43. A recent study reported that RNF126 targets aggregation-prone fragments of TDP43 for proteasomal degradation in concert with BAG6.^52^

**Figure 6.**
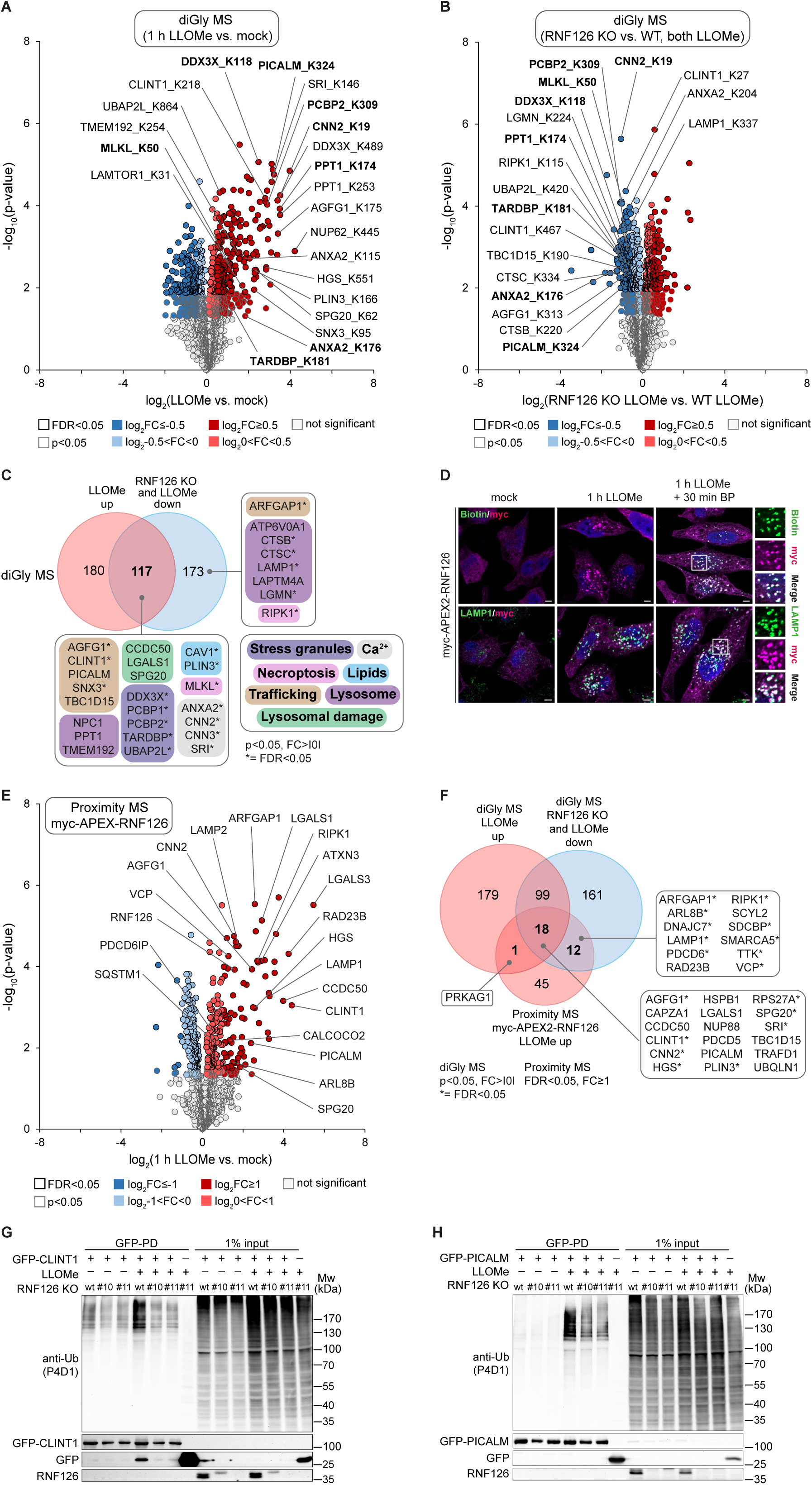
RNF126 substrates possess diverse functions associated with damaged lysosomes (A and B) Volcano plots of altered ubiquitylation sites in RNF126 WT cells treated with LLOMe or left untreated (A) and in LLOMe-treated RNF126 WT or KO cells (B) identified by diGly MS. Significantly altered proteins are labeled in dark (log_2_FC ≥ I0.5I) and light color (log_2_I0<FC<0.5I), respectively. Among these, black circles indicate a significance of FDR<0.05, whereas grey circles a threshold of p<0.05. Damage-induced and RNF126 sensitive ubiquitylation sites in selected proteins are highlighted. Selected ubiquitylation sites that increased in (A) and decreased in (B) are marked in bold. (C) Potential damage-induced RNF126 substrates from the overlap of significantly regulated diGly sites (p<0.05, log_2_FC>I0I) increased in (A) and decreased in (B) are annotated manually. Hits reaching a significance threshold of FDR<0.05 are marked with *. (D) myc-APEX2-RNF126 expressing cells treated as indicated were fixed and stained with indicated antibodies. Insets show magnifications of boxed areas. Scale bars, 5 µm. (E) Volcano plot of changes in the proximity proteome of RNF126 upon LLOMe. Experiments were performed in biological quadruplicates and statistical analysis using two-sided two sample Student’s t-test (p-value<0.05) with permutations-based correction (FDR<0.05). Significantly altered proteins are labeled in dark (log_2_FC ≥ I1I) and light color (log_2_I0<FC<1I), respectively. Among these, black circles indicate a significance of FDR<0.05, whereas grey circles a threshold of p<0.05. Selected proximity partners are highlighted. (F) Venn diagram of proteins that showed increased ubiquitylation (p<0.05, log_2_FC≥0) upon damage, were found close to RNF126 upon damage (FDR<0.05, FC ≥ 1) and showed reduced ubiquitylation in damaged RNF126 KO cells (p<0.05, log_2_FC<0). Overlapping hits are considered potential damage-induced RNF126 substrates. Proteins marked with * reach a significance threshold of FDR<0.05 in both diGly data sets. (G and H) GFP-CLINT1 (G) and GFP-PICALM (H) expressed in RNF126 WT and KO cells (two independent clones) were subjected to denaturing GFP-IP and immunoblotting.

To complement our diGly proteomics and to stringently narrow down RNF126’s substrate spectrum, we performed APEX2-based proximity labeling using cells expressing myc-APEX2-RNF126 to capture RNF126 substrates by an independent approach. We confirmed proper expression of the APEX2 fusion construct, its biotinylation capacity and successful translocation to damaged lysosomes (Figures 6D, S5D, S5E). Statistical analysis of RNF126’s protein proximity upon LLOMe revealed a clear lysosome signature with enrichment of lysosomal proteins (LAMP1, LAMP2), damage markers (SPG20, LGALS1, LGALS3) and the lysophagy receptor CCDC50.^53^ Similarly to our TMEM192-APEX2 proximity labeling (Figure 1B), we also observed an accumulation of ESCRT members (HGS, PDCD6IP) and other autophagy components (p62, CALCOCO2 (also known as NDP52)). Importantly, we detected several ubiquitylation targets from our diGly MS approach in close vicinity to RNF126 (Figure 6E, Table S5). By systematically comparing diGly and RNF126 proximity data, we found an overlap of 18 significantly regulated proteins (Figures 6F) which we considered candidate RNF126 substrates. Among them were several established lysosomal damage response components (CNN2, CCDC50, HSPB1, LGALS1, SPG20, TBC1D15) as well as a number of endocytic proteins (PICALM, CLINT1, AGFG1 (also known as HRB)) that are known to be ubiquitylated in response to lysosomal damage.^21^ From the latter category, we biochemically validated RNF126- and LLOMe-dependent ubiquitylation of PICALM and CLINT1 using denaturing pulldowns (Figures 6G, 6H), implicating that both RNF126 substrates are indeed part of the lysosomal damage response. However, since RNF126 deficiency did not affect the abundance of CLINT1 or PICALM1 (Figure S5F), it seems that RNF126-mediated ubiquitylation exert non-degradative, regulatory functions on these substrates. Collectively, these results show that RNF126 ubiquitylates known and potential new components of the lysosomal damage response and is therefore an important part of this pathway.

### RNF126 deletion affects regeneration of damage lysosomes

Next, we asked whether and how RNF126 deficiency impacts on the lysosomal damage response pathway. Firstly, we monitored Gal3 accumulation in time course and pulse-chase experiments in RNF126 WT and KO cells. Interestingly, the latter exhibited increased Gal3 punctae starting at 30 min post-LLOMe treatment as well as elevated numbers of persistent Gal3-positive structures up to 2 h after LLOMe washout (Figures 7A, S6A). However, since RNF126 KO cells were able to clear Gal3 punctae at the end of our 3 h-time course (Figure S6A) and also did not show a reduction in Gal3-based lysosomal flux (Figure S6B), we concluded that RNF126 is dispensable for lysophagy.

**Figure 7.**
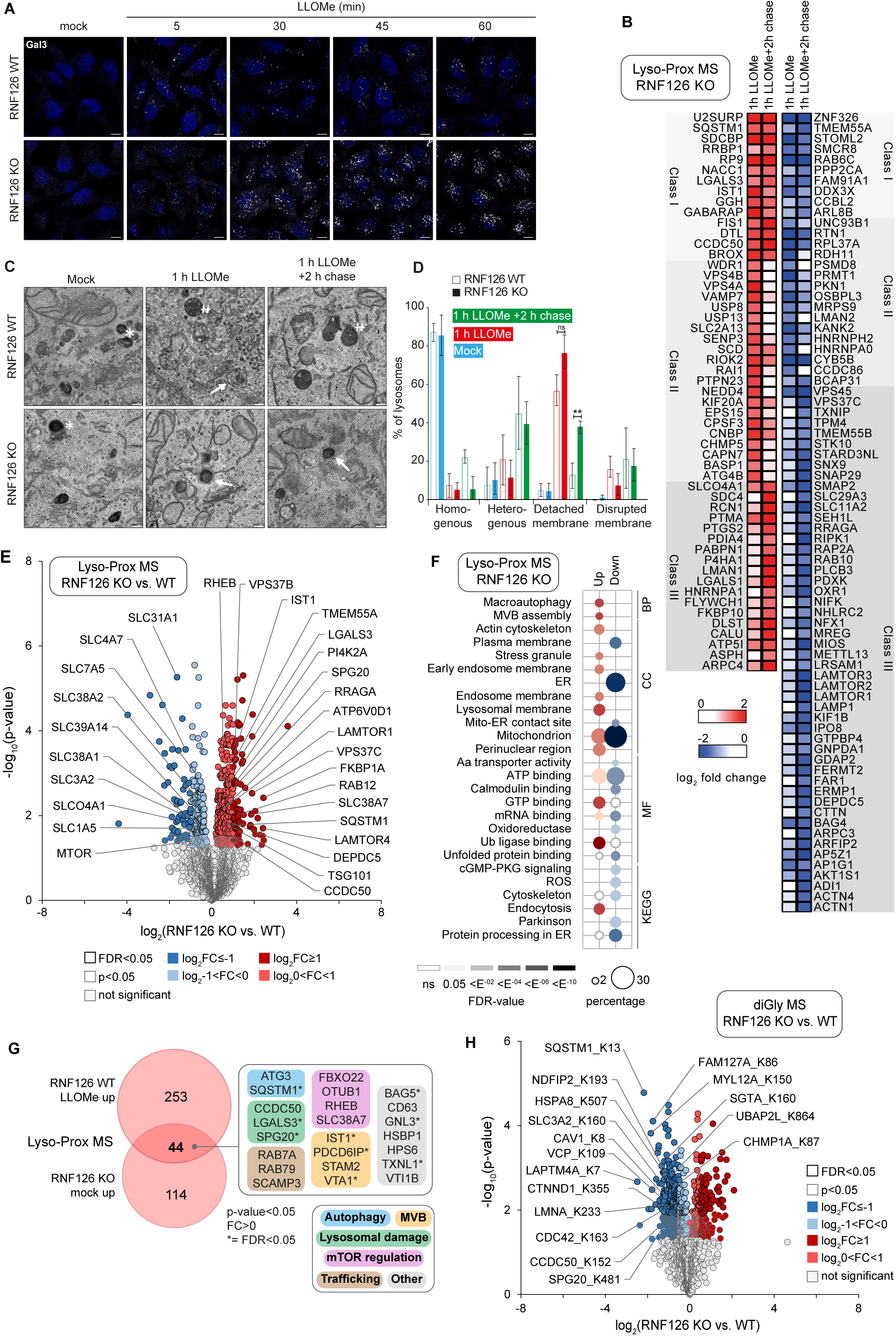
RNF126 functions in regeneration damage response and lysosomal homeostasis (A) RNF126 WT and KO cells were treated up to 60 min with LLOMe, fixed and stained with an anti-Gal3 antibody. Scale bars, 10 µm. (B) Heat map of Lyso-Prox MS in RNF126 KO cells treated with LLOMe as indicated or left untreated. TMEM192 proximity partners that decrease or increase in abundance at lysosomes upon 1 h LLOMe or 1 h LLOMe followed by 2 h chase compared to mock treatment. Based on p<0.05 and LLOMe-induced abundance pattern, proteins were grouped in three classes. Class I consists of proteins whose abundance consistently changed irrespective of the LLOMe treatment whereas Class II and III contains proteins whose abundance only changed upon 1 h LLOMe or 1 h LLOMe followed by 2 h chase, respectively. (C) TEM of lysosomes in parental and RNF126 KO cells treated as indicated. Arrows indicate lysosomes with swollen limiting membrane detached from the lipid-dense core. * labels homogenous lysosomes. # represents heterogenous lysosomes with enlarged dense core. Scale bars, 200nm. (D) Quantification of lysosomal morphology classes described in (G). The plot represents mean and standard deviation from three biological replicates. **p < 0.01. (E) Volcano plot of TMEM192-Flag-APEX2 proximity proteome changes between RNF126 WT and KO cells. Experiments were performed in biological quadruplicates and statistical analysis using two-sided two sample Student’s t-test (p-value<0.05) with permutations-based correction (FDR<0.05). Significantly altered proteins are labeled in dark (log_2_FC ≥ I1I) and light color (log_2_I0<FC<1I), respectively. Among them, black circles indicate significance of FDR<0.05, whereas grey circles a threshold of p<0.05. (F) GO enrichment analysis of proteins significantly enriched and diminished (FDR<0.05, log_2_FC>I0I) at lysosomes from (E). Dot size correlates to number of proteins, dot color to term enrichment (FDR). BP, biological process; CC, cellular component; MF, molecular function; KEGG, Kyoto Encyclopedia of Genes and Genomes. (G) Venn diagram showing overlap of proteins enriched at lysosomes (p<0.05, log_2_FC>0) in LLOMe-treated RNF126 WT cells -and mock-treated RNF126 KO cells (E). Selected proteins were annotated manually. Overlapping proteins with a significance threshold of FDR<0.05 are marked with (*). (H) diGly MS of RNF126 WT and KO cells in untreated condition. Significantly altered proteins are labeled in dark (log_2_FC ≥ I0.5I) and light color (log_2_I0<FC<0.5I), respectively. Among these, black circles indicate a significance of FDR<0.05, whereas grey circles a threshold of p<0.05. See also Figures S6 and S7.

Secondly, we used lysosomal proximity labeling coupled to quantitative mass spectrometry to unbiasedly map lysosomal proteome changes in RNF126 KO cells comparing mock treatment to 1 h LLOMe or 1 h LLOMe followed by 2 h washout with fresh media. Importantly, expression of TMEM192-Flag-APEX2, its biotinylation capacity and lysosomal localization of biotin did not differ between RNF126 WT and KO cells (Figures S6C, S6D). Comparative analysis of lysosome proximal proteins with significantly altered abundance across both data sets (Figures S6E, S6F, Table S6) revealed three classes of proteins that differed in their response to damage when RNF126 was absent (Figure 7B). Class I consists of proteins with consistent alterations in both LLOMe treatments. Among Class I proteins were several components of the lysosomal damage response (LGALS3, GABARAP, p62, IST1, CCDC50) which were enriched as well as the members of the GTPase system (ARL8B, RAB6C, SMCR8) and the phosphatidylinositol phosphatase TMEM55A (also known as PIP4P2) that were decreased on damaged lysosomes (Figure 7B). The other two classes contain proteins responsive to either LLOMe (Class II) or LLOMe followed by a recovery phase (Class III). Intriguingly, among Class II proteins that acutely increased in their abundance were ESCRT members (CHMP5, VPS4A, VPS4B, PTPN23), endocytic proteins (NEDD4, EPS15, USP8, VAMP7) and autophagy regulators (ATG4B, USP13) while Class III proteins which decreased in abundance in the LLOMe recovery phase consists of numerous mTORC1 components (LAMTOR1, LAMTOR2, LAMTOR3, DEPDC5, AKT1S1, RRAGA) and functionally diverse lysosomal regulators (RAB10, TMEM55B, SNAP29, LAMP1, SLC29A3, SLC11A2, KIF1B) (Figure 7B). Overall, this analysis shows that lysosomes from cells lacking RNF126 are still able to respond to damage in a similar manner than those from LLOMe-treated parental cells (Figures S7A). However, a number of proteins critically involved in lysosomal homeostasis (SMCR8, ARFGAP1), vesicle trafficking (RAB6C, ARFIP2) and endocytosis (SNX9, CLINT1) exclusively lost their lysosomal proximity upon damage in a RNF126-dependent manner (Figure S7A). For the *bona fide* RNF126 substrate CLINT1 (Figure 6G) which followed a similar kinetic as RNF126 with regard to recruitment to and persistence on damaged lysosomes (Figures 3A, 3D), we confirmed reduced lysosomal localization in the absence of RNF126 in LLOMe-treated cells (Figures S7B). Given that RNF126 arrives after initial damage detection but does not impair lysophagy and that CLINT1 has also not been associated with lysosomal repair or lysophagy, we propose that RNF126-mediated ubiquitylation of a set of functionally diverse substrates (among them CLINT1) contributes to the regeneration phase of the lysosomal damage response in which lysosomes have undergone initial repair and triaged away from lysophagy but did not yet regain full functionality as they remain non-acidified.

To test this hypothesis, we examined the morphology of lysosomes in RNF126 WT and KO cells using transmission electron microscopy. In the absence of damage, lysosomes did not show any overt changes at the ultrastructural level (Figures 7C, 7D). Upon acute LLOMe treatment lysosomes shifted from an overall homogenous appearance to three morphologically distinct sub-populations characterized by a heterogenous appearance, a dense core with a detached limiting membrane and a disrupted membrane. Even though no substantial changes in the proportional distribution of these sub-populations upon RNF126 deficiency were observed after 1 h LLOMe treatment, a significant portion of lysosomes from RNF126 KO failed to recover from LLOMe treatment and showed persistent detachment of the limiting membrane after 2 h chase (Figures 7C, 7D). Of note, this defect did not result in a volume increase of the recovering lysosomes (Figures S7C). As a consequence of this lysosomal regeneration defect, loss of RNF126 shifted the lysosomal population to undergo even more lysophagy based on flux analysis (Figure S6B). Together, these findings support the notion that RNF126 is required for the regeneration phase of the lysosomal damage response.

### RNF126 also helps to maintain lysosomal homeostasis

Intrigued by the elevated Gal3 response to lysosomal damage in cells lacking RNF126 (Figure 6A), we asked whether this may reflect a predisposition to lysosomal damage. This would imply that RNF126 might be required for lysosomal homeostasis besides its function in response to lysosomal damage. To assess whether lysosomes in RNF126 KO cells exhibit damage features under basal growth conditions compared to their wild-type counterparts, we performed lysosomal proximity profiling using TMEM192-Flag-APEX2 expressing RNF126 WT and KO cells (Figure S6C, S6D).

In agreement with our hypothesis, lysosomes of RNF126 KO cells showed signs of damage in the absence of LLOMe treatment, exemplified by accumulation of damage markers (LGALS3, SPG20, CCDC50, RAB12), ESCRT proteins (IST1, TSG101, PDCD6IP, VPS37B, VPS37C) and the PITT pathway component PI4K2A (Figure 7E, Table S6). Consistent with that, GO term analysis revealed an accumulation of autophagy machinery and MVB components, stress granule and actin cytoskeleton proteins as well as proteins involved in Ub ligase binding and located in the perinuclear region (Figure 7F), all reflecting fragility of lysosomes upon RNF126 loss. Moreover, there was considerable overlap of proteins enriched on lysosomes from LLOMe-treated RNF126 WT cells and mock-treated RNF126 KO cells (Figure 7G, Table S6). In line with an unbalanced lysosomal homeostasis, we also found that mTORC1 complex components (RHEB, LAMTOR1, LAMTOR4, DEPDC5, PRAGA, SLC38A7, FKB1A) were enriched on lysosomes in RNF126 KO cells whereas mTOR itself was found increasingly dissociated. Additionally, we found reduced lysosomal association of numerous solute carrier (SLC) transporters involved in ion and amino acid transport (Figure 7E, Table S6). GO term analysis confirmed a loss of plasma membrane proteins and amino acid transporters next to mitochondrial and ER proteins in the proximity of lysosomes in RNF126 KO cells (Figure 7F). Hence, RNF126 plays a key role in maintaining lysosomal homeostasis.

To better understand how RNF126 contributes to lysosomal homeostasis we turned again to diGly proteomics to this time identify constitutive RNF126 substrates in basal growth conditions (Figure S7D). While overall protein abundance remained unaffected in RNF126 KO cells (Figure S7E), several proteins showed reduced ubiquitylation when RNF126 was absent (Figure 7H, Table S7). Among these were the lysosomal damage components SPG20, CCDC50 and p62 as well as the transmembrane proteins SLC3A2 and LAPTM4A of which the latter exhibited the strongest RNF126-regulated ubiquitylation site (Figure 7H). Intriguingly, members of the LAPTM family are known to be sorted from the plasma membrane to lysosomes in an Ub-dependent manner.^54^ Since RNF126 has been implicated in Ub-mediated sorting of epidermal growth factor receptor (EGFR) and cation-independent mannose 6-phosphate receptor (CI-M6PR)^55, 56^, it is plausible that RNF126 exerts a similar function for LAPTM4A. In turn, LAPTM4B, a homologue of LAPTM4A, has been reported to interact with the heterodimer SLC3A2/SLC7A5 at the plasma membrane and to mediate its transport to lysosomes where SLC3A2/SLC7A5 promotes leucine influx resulting in mTORC1 activation mediated via v-ATPase stimulation.^57^ Hence, the altered lysosomal abundance of several SLC transporters as well as mTORC1 and v-ATPase complex components (Figure 7E) might be a consequence of defective sorting of LAPTM4A/B and SLC3A2/SLC7A5 in the absence of RNF126-mediated ubiquitylation of these targets. Collectively, we propose that besides driving regeneration from lysosomal damage RNF126 exerts a housekeeping role in maintaining lysosomal homeostasis in unstressed condition.

## DISCUSSION

In this study, we used proximity labeling and organellar proteomics to profile LLOMe-induced changes at lysosomes and identified the RING E3 ligase RNF126 as novel regulator of ubiquitylation events on damaged lysosomes. We observed translocation of RNF126 to lysosomes upon various lysosomal insults independent of Gal3 and ATG8-ylation of the lysosomal membrane and after the initial ESCRT-mediated repair phase. RNF126 depletion resulted in strong reduction of LLOMe-induced ubiquitylation of lysosomal but also non-lysosomal proteins. However, RNF126 loss did not lead to compromised lysosomal clearance via lysophagy, suggesting that RNF126-catalyzed ubiquitylation is not primarily serving as degradative signal for lysophagy but also exhibits regulatory functions.

One prominent category of LLOMe-regulated and RNF126-dependent ubiquitylated proteins represented those involved in stress granule (SG) formation (DDX3X, UBAP2L, TDP-43).^58^ Rapid SG formation was observed upon lysosomal damage in previous studies.^50, 51^ Interaction of SG proteins NUFIP2 and G3BP1 with the lysosomal membrane was shown to be mediated by non-canonically lipidated ATG8. Since NUFIP2 was found to negatively regulate mTOR activity,^50^ SG formation may couple storage of translation components and shutdown of mTOR-mediated translation as possible way to ensure a coordinated response to damage. Which role ubiquitylation might play on SG assembly, disassembly and association with the lysosome in response to damage is currently unknown and requires further investigation.

Another category of RNF126 substrate candidates comprised proteins involved in endocytic pathway and vesicular transport. RNF126 ubiquitin ligase activity was previously shown to be crucially involved in transmembrane trafficking, including retrograde sorting of CI-M6PR, a transporter for lysosomal hydrolases, providing first evidence for a role of RNF126 in regulating lysosomal function.^56^ Depletion of RNF126 retains CI-M6PR in endosomes and increases lysosomal degradation of the receptor. However, Smith and McGlade were unable to identify CI-M6PR as a direct RNF126 substrate.^56^ Ubiquitylation of the endocytic machinery instead of the cargo was proposed as alternative mechanism to finetune protein sorting. It was shown previously that the clathrin associating phosphoinositide binding protein CLINT1 (epsinR) is involved in retrograde sorting of M6PR.^56, 59^ Next to its role in retrograde trafficking, the N-terminal ENTH domain (with high affinity for PI4P)^60^ of CLINT1 was shown to bind to SNARE protein VTI1B^61^ that functions in homotypic late endosome fusion together with STX8, STX7 and VAMP8.^62^ Strikingly, we found ubiquitinated CLINT1 accumulated on lysosomes upon damage in a RNF126-dependent manner. Since we detected multiple damage-induced ubiquitylation sites on CLINT1 (K6, K27 K183, K192, K218, K260, K283, K314, K467), its ubiquitin-mediated regulation seems multilayered. Interestingly, two of these ubiquitylation sites (K6 and K27) reside within the ENTH domain responsible for SNARE binding. We therefore hypothesize that ubiquitylation might serve as a block to prevent fusion events with compromised lysosomes. In contrast to CI-M6PR, EGFR was shown to be directly ubiquitylated by RNF126. The N-terminal Ub-binding zinc finger motif was hereby necessary for the association with the transmembrane receptor.^55^ Additionally, RNF126 as well as its homologue RNF115 (Rabring7) were shown to be involved in endosomal sorting of other transmembrane proteins, including MET, CXCR4 and MET.^55^ Presumably, ubiquitylation substrates in context of lysosomal damage might also interact with the N-terminal Zinc finger domain of RNF126.

In addition to Ub-mediated sorting processes, RNF126 is implicated in cytosolic protein quality control of mislocalized membrane proteins in the cytosol.^47, 48^ In this context, substrate specificity was mediated by chaperone BAG6 that interacts with the RNF126’s N-terminal Zinc finger motif.^47, 49^ Despite its importance in the aforementioned pathways, however, the Zinc finger motif was neglectable for the recruitment of RNF126 to damaged lysosomes. Instead, we found an arginine-stretch in a low complexity region without prominent structural features. Charge-reversing mutation abolished the recruitment completely. Previous damage-responsive E3 ligases were either shown to be directly recruited to the lysosome for examples by binding to exposed glycans (SCF^FBXO^^27^)^19^ or interaction with lysosomal membrane proteins such as LAMP2 (CUL4A-DDB1-WDFY1)^20^ or employ an soluble adaptor that facilitate lysosomal recruitment. Examples for this latter scenario are TRIM16 and ITCH which are recruited by Gal3 and SPG20, respectively.^33, 33^ Intriguingly, SPG20 and RNF126 labeled very similar lysosomal populations and both responded to lipid peroxidation inducing agents. Since peroxidized lipids exert additional negative charges, we hypothesize that the arginine-rich stretch of RNF126 undergoes a direct electrostatic interaction with peroxidized lipids. However, we cannot exclude that lysosomal recruitment of RNF126 involves an adaptor.

RNF126 can catalyze several types of Ub chains including K48 and K63 Ub chains. Both are induced upon lysosomal damage. While K63 Ub was described as recruitment signal for Ub-binding receptors to facilitate phagophore formation around the damaged lysosome, K48 Ub on lysosomal transmembrane proteins was shown to sterically block autophagosome maturation and subsequently be extracted by the triple ATPase VCP.^18, 31^ We found a strong reduction in K48 Ub at the lysosome. Further experiments are necessary to examine whether lysosomal K63 Ub is also mediated by RNF126.

Lastly, we uncovered a potential constitutive role of RNF126 in regulating lysosomal homeostasis. Next to regulating endosomal transport of surface membrane receptors, we identified the lysosomal membrane protein LAPTM4A as RNF126 substrate. LAPTM4A was identified as short-lived lysosomal transmembrane protein being degraded by internalization into the lysosomal lumen via the ESCRT machinery. This process was previously described to be mediated by NEDD4.^34^ In contrast to NEDD4 that binds to carboxyl-terminal PY motifs^34, 57^, RNF126-dependent ubiquitylation was detected at the cytosol-facing N-terminus (lysine 7 (K7)). We are currently elucidating how RNF126 contributes to sorting, degradation or interaction of LAPTM4A with other factors. For instance, we also found the SLC transporter SLC3A2 to be less ubiquitylated in RNF126 KO cells. Interestingly, LAPTM4B, a homologue of LAPTM4A, has been reported to interact with the heterodimer SLC3A2/SLC7A5 at the plasma membrane and to mediate its transport to lysosomes.^57^ SLC1A5, SLC3A2, and SLC7A5 are majorly implicated in the uptake of glutamine and leucine^63^ and where all found reduced in abundance at the lysosome in the absence of RNF126. Leucine influx at the lysosome was shown to activate mTORC1.^57^ Interestingly, we found mTOR dissociated from lysosomes in RNF126 KO cells as well as reduced abundance of several SLC transporters including SLC1A5, SLC3A2, and SLC7A5, pointing towards a regulatory function of RNF126 in nutrient sensing at the lysosome. To sum up, we find multiple pathways that are regulated by RNF126 ubiquitylation converging at the lysosome and affecting lysosomes in stressed as well as unstressed conditions.

## ACKNOWLEDGMENTS

We are thankful to members of the Behrends lab for critical discussions and shared reagents. We thank Harald Stenmark, Vojo Deretic and Sankarshana Nagaraja for providing PI4K2A KO, GAL3 KO and RNF126 WT/KO TMEM192-FLAG-APEX2 cells, respectively. This work was supported by the Deutsche Forschungsgemeinschaft (DFG, German Research Foundation) within the frameworks of the Munich Cluster for Systems Neurology (EXC 2145 SyNergy, Project-ID 390857198), the Collaborative Research Center 1177 (ID 259130777) and the project grant BE 4685/7-2 (ID 447112704). The latter is a joint grant awarded to C.B. and H.M..

## AUTHOR CONTRIBUTIONS

S.L. performed all experiments except nano-silica and laser irradiation (B.K.), denaturing pulldowns (B.K., P.G), microglia preparations (L.C.), initial Lyso-IP and diGly MS (L.C., H.T.), EM (M.S., G.K.), DNA-PAINT (K.C., R.J.), RNF126 antibody production (R.F.) and generation of Cd68-Flag knock-in mice (B.W., W.W.). All authors including H.M. participated in data analysis and discussion. S.L. and C.B. conceived the study and wrote the manuscript.

## STAR METHODS

### KEY RESOURCES TABLE

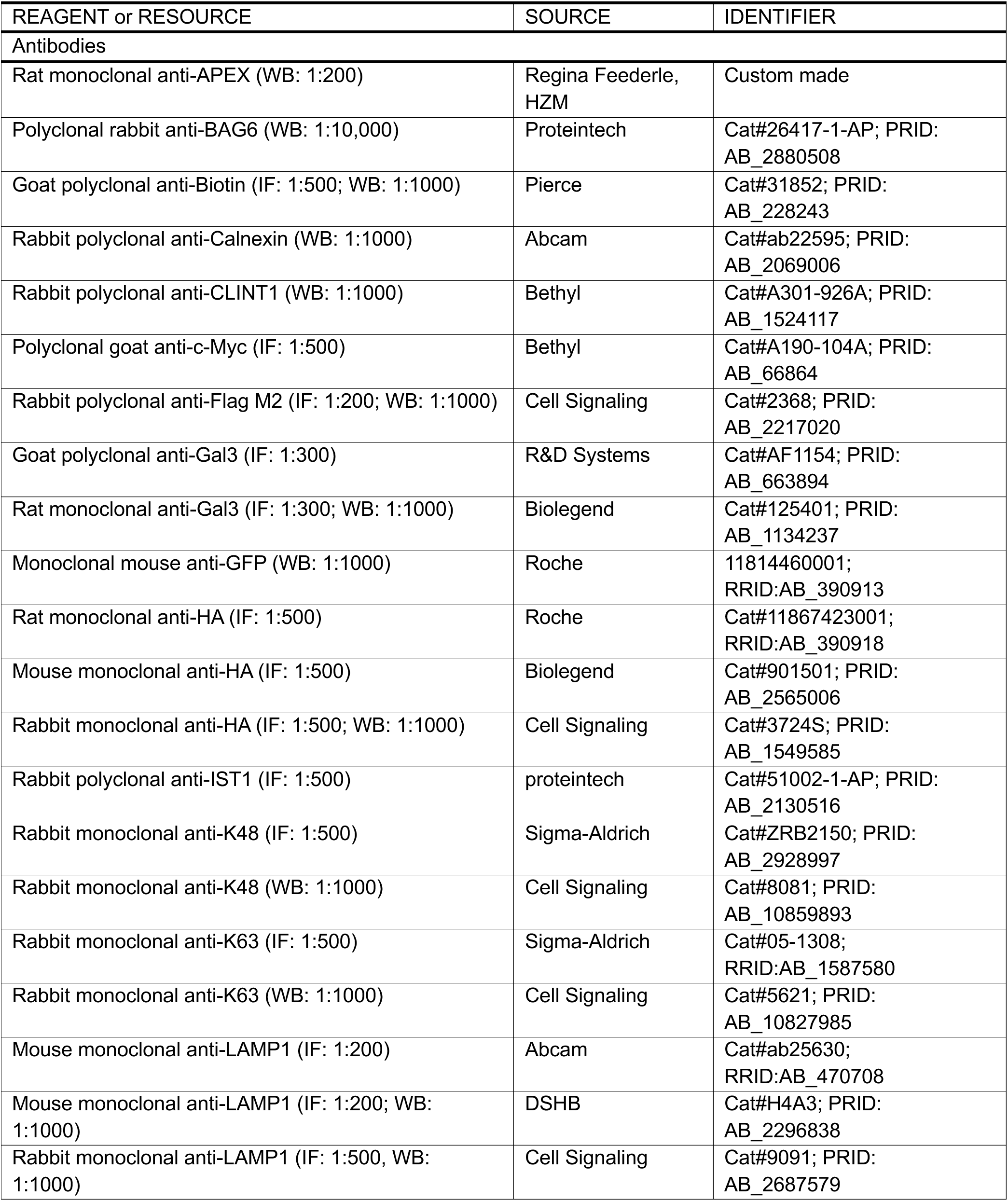
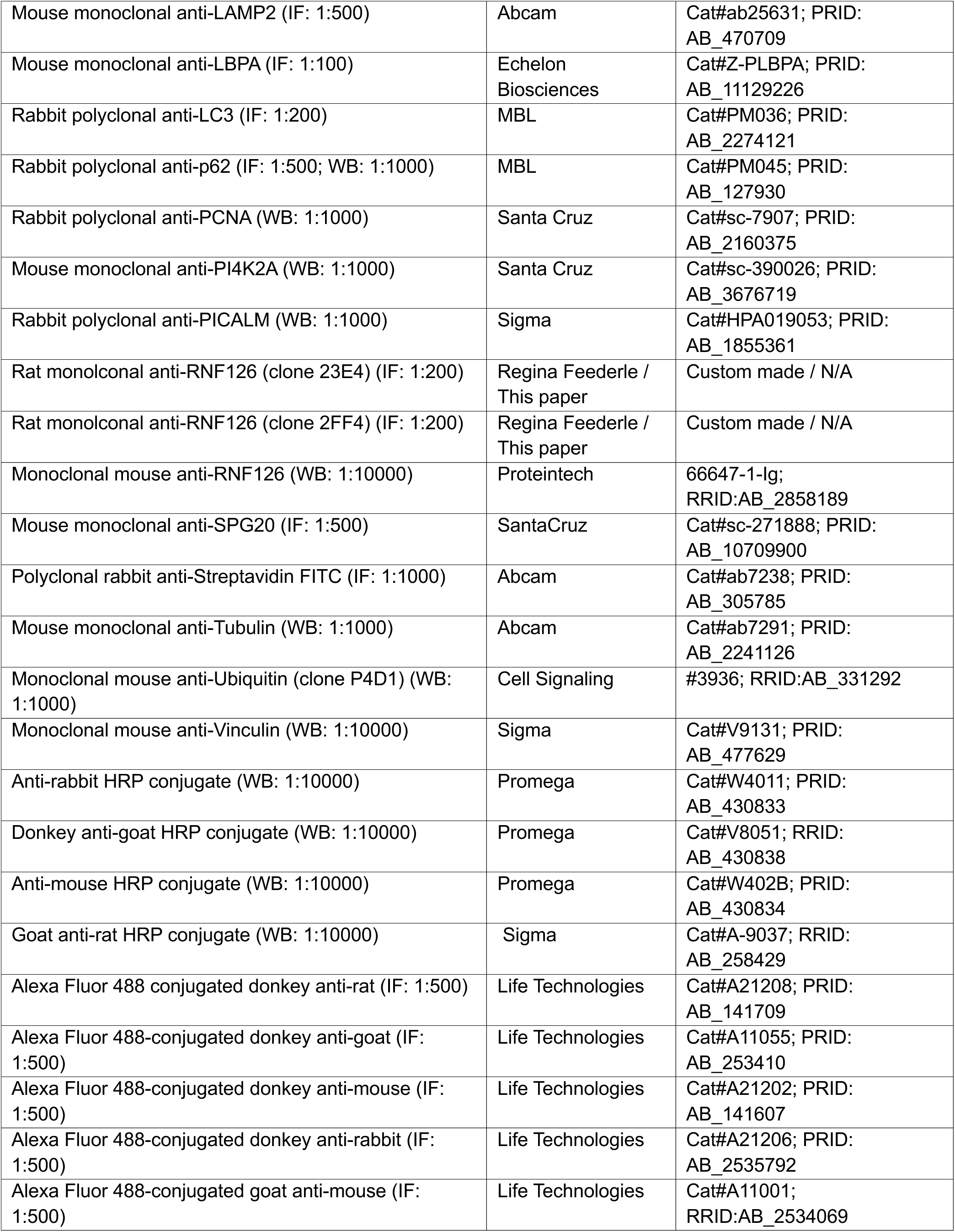
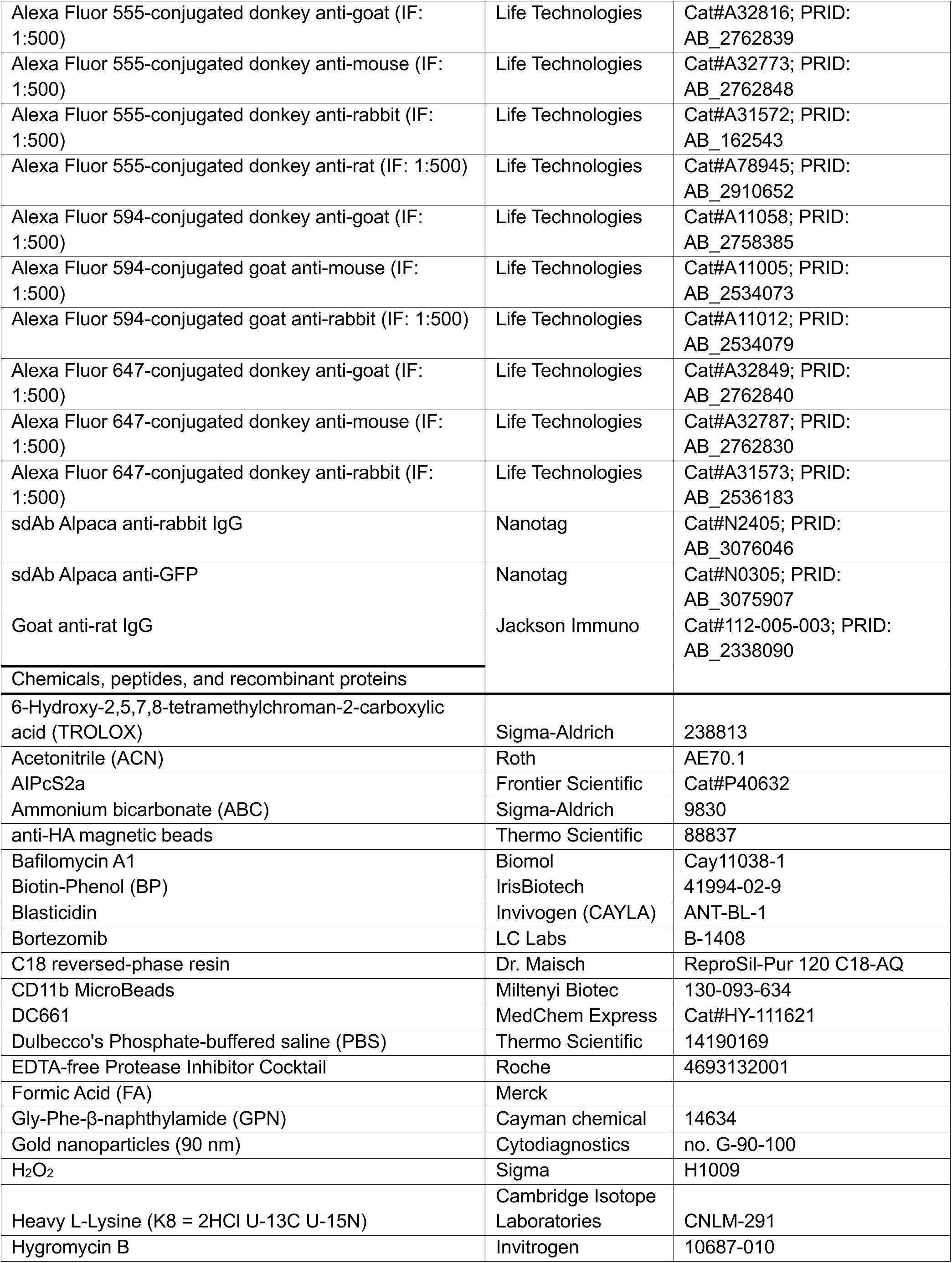
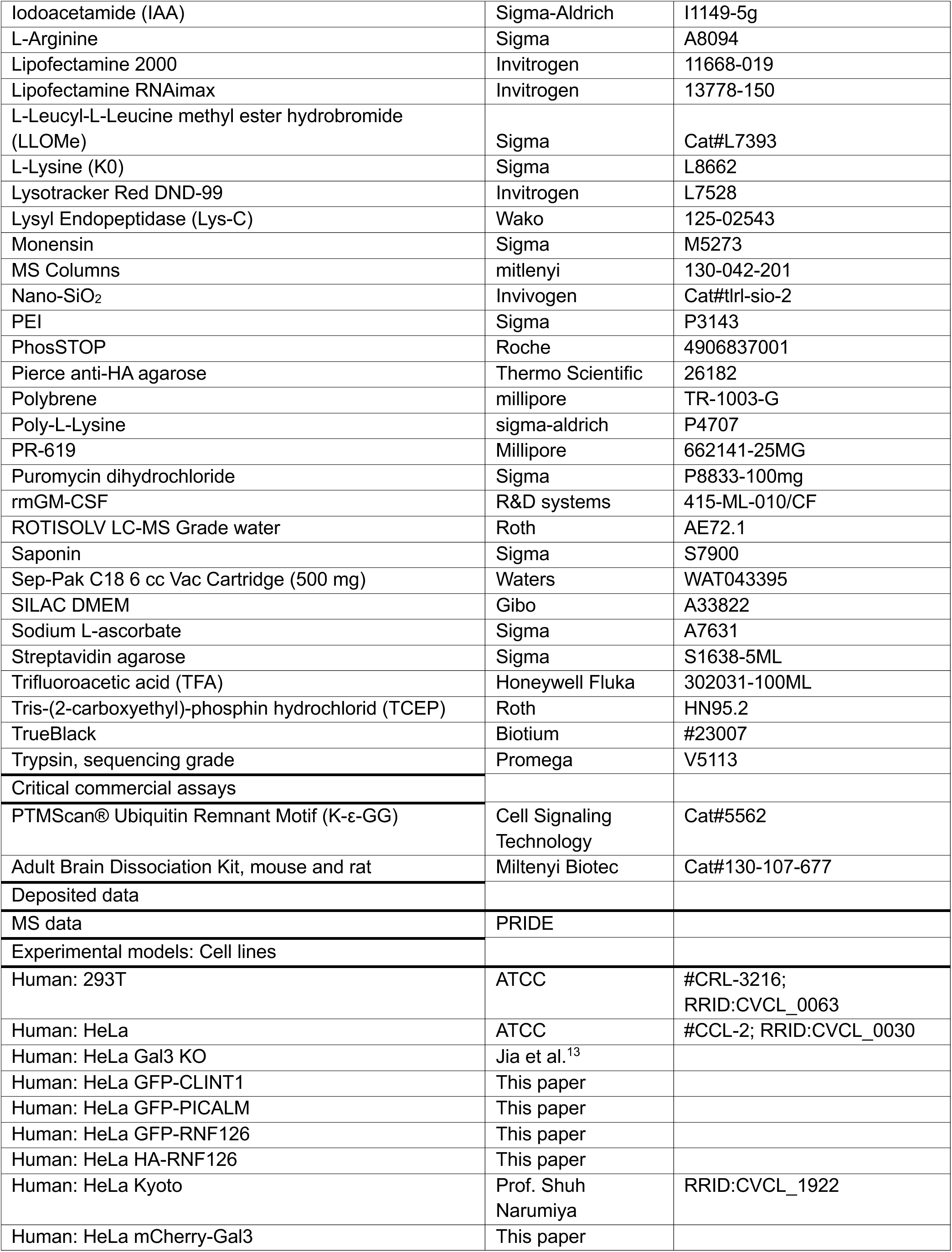
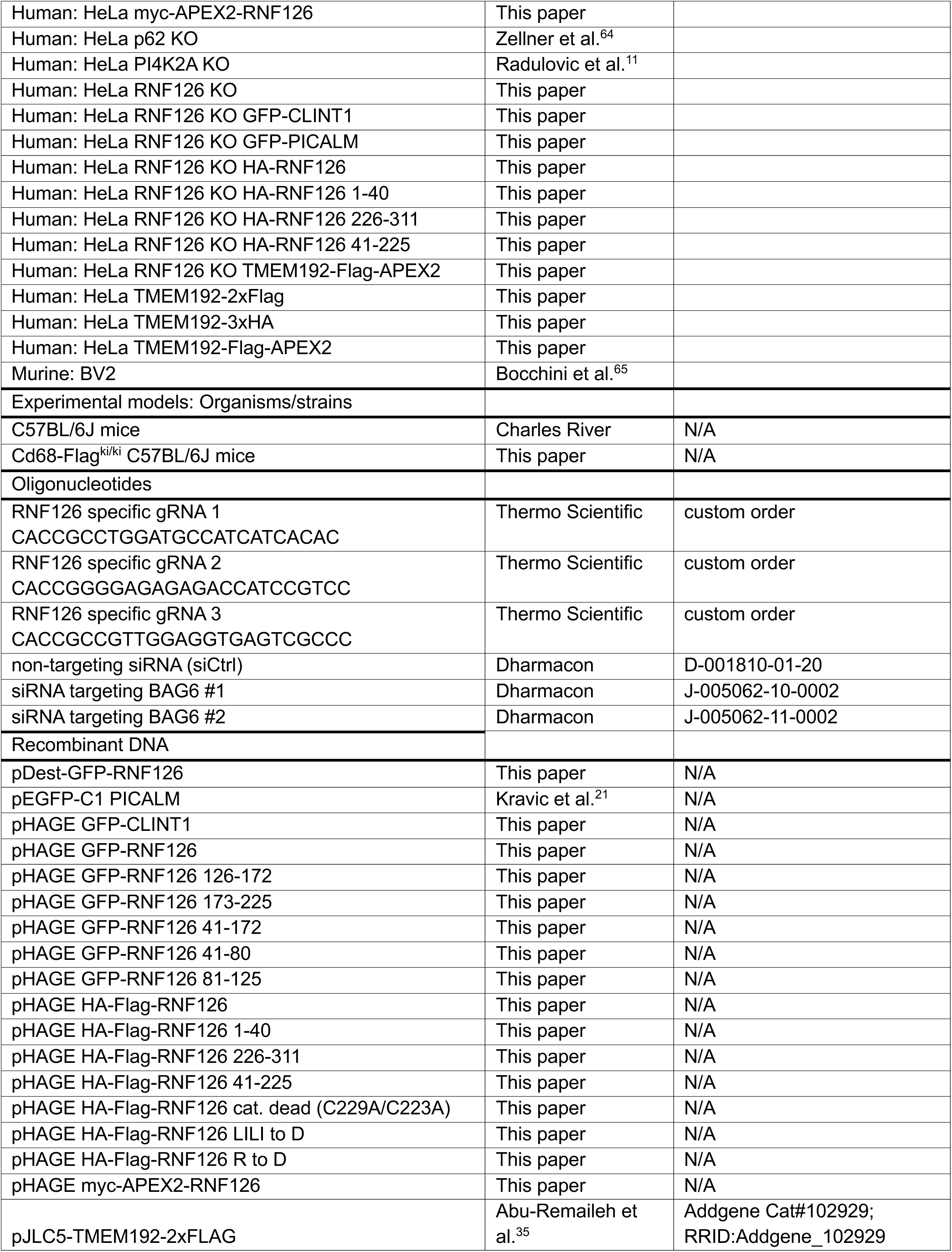
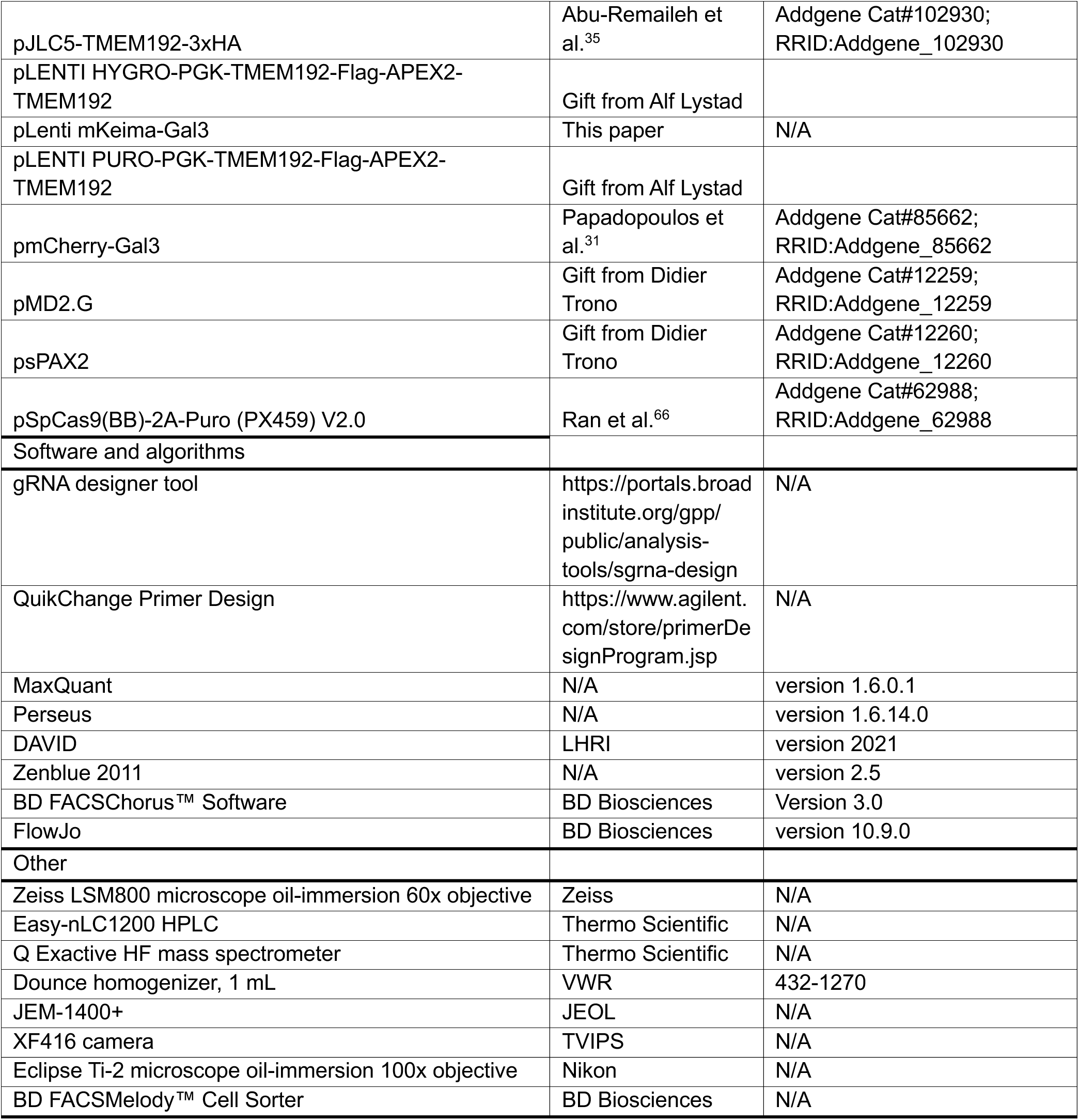

### RESOURCE AVAILABILITY

#### Lead Contact

Further information and requests for resources and reagents should be directed to and will be fulfilled by the Lead Contact, Christian Behrends.

#### Materials Availability

All unique reagents generated in this study are available from the Lead Contact with a completed Materials Transfer Agreement.

### METHODS

#### Cloning

For plasmids, the Gateway® system was used. ORFs were flanked with attB sites using PCR, transferred into the entry vector pDONR223 and introduced into destination vectors via recombination reactions. For CRISPR/Cas9-mediated knockout cell line generation, sgRNAs were picked with the Broad Institute GPP sgRNA designer tool (https://portals.broadinstitute.org/gpp/public/analysis-tools/sgrna-design) and ligated with the Cas9 encoding BbsI-digested pSpCas9(BB)-2A-Puro (PX459) V2.0 vector (Addgene, #62988). To introduce point mutations, mutagenesis primers were designed with the QuikChange Primer Design Software tool (Agilent Technologies). Forward and reverse primers were used in individual PCRs using KOD Hot Start polymerase (Merck Millipore) according to manufacturer’s instructions. PCRs were then combined followed by a second round of PCR. After DpnI digest of template DNA, PCR products were purified using the QIAquick PCR purification kit (Qiagen) according to manufacturer’s instructions and transformed in XL10 Gold *E. coli*. All cloning products were verified by Sanger Sequencing.

#### Cell Culture

HeLa (ATCC), 293T (ATCC) and murine microglial cell line BV2^65^ were cultured in Dulbecco’s modified Eagle’s medium (DMEM) (Gibco) supplemented with 10% fetal bovine serum (FBS) and 1 mM sodium pyruvate (Gibco) and grown at 37°C and 5% CO_2_. For stable overexpression cell lines antibiotics were added. For SILAC diGly-MS experiments, HeLa cells were grown in lysine and arginine-free SILAC DMEM (Gibco) supplemented with 10% dialyzed FBS, 2 mM L-glutamine (Gibco), 1 mM sodium pyruvate (Gibco), 38 µg/ml arginine (Sigma) and 66 µg/ml light (K0, Sigma) or heavy L-lysine (K8 = 2HCl U-13C U-15N, Cambridge Isotope Laboratories). Primary microglia isolated from adult mice were kept in DMEM/F-12 (Gibco), supplemented with 10% FBS, 1 mM sodium pyruvate, 1X MEM NEAA (Gibco), 1% Pen/Strep (Gibco) and 10 ng/ml recombinant mouse GM-CSF (R&D Systems). HeLa cells were used if not stated otherwise.

#### Cell line generation

Stable overexpression cells were obtained by lentiviral transduction followed by antibiotic selection. Briefly, 293T cells were transfected with 2 µg lentiviral pHAGE constructs and packaging plasmids (2.7 µg psPAX2 (Addgene, #12260), 1 µg pMD2.G (Addgene, #12259)) using 10 µl Lipofectamine 2000 (Thermo Fisher Scientific). Virus-containing media was collected after 24 h and 48 h, combined, filtered, mixed with 8 µg/ml polybrene (Millipore) and added on recipient cells. Next day, media was replaced. 24 h later, media was changed to media containing 2 µg/ml puromycin (Sigma), 4 µg/ml blasticidin (Invivogen), or 200 µg/ml hygromycin B (Invitrogen). CRISPR/Cas9-mediated genome editing was used to generate RNF126 KO cells. A pool of 3 individual sgRNAs targeting different loci in the RNF126 gene were introduced in HeLa cells via transfection with Lipofectamine 2000 in a 1:3 ratio (µg DNA: µl Lipofectamine). 24 h post-transfection, cells were selected with 2 µg/ml puromycin for 48 h. Afterwards, cells were kept in normal growth media and single cell clones were obtained via serial dilution. Single cell KO clones were validated via Sanger sequencing of genomic DNA and immunoblotting.

#### Generation of Cd68-Flag mice

Cd68-Flag mice were generated by CRISPR/Cas9-assisted gene editing in zygotes as described previously.^67^ Briefly, pronuclear stage zygotes were obtained by mating C57BL/6J males with superovulated C57BL/6J females (Charles River). Embryos were then electroporated using the NEPA21 electroporator and a 1 mm electrode with a CD68-specific CRISPR/Cas9 ribonucleoprotein (RNP) solution consisting of 200 ng/µl SpCas9 WT protein (IDT), 6 µM crRNA (protospacer GTAGGTTGATTGTCGTCTGC; IDT), 6 µM tracrRNA (IDT), and 33 ng/μl of a mutagenic 478 bp single-stranded oligodeoxynucleotide for the introduction of a 3xFLAG tag at the C-terminus. The lengths of the homology arms were 200 nt each. After microinjection, treated zygotes were transferred into pseudopregnant CD-1 foster animals. Mutant founder animals were crossed to C57BL/6J animals to isolate the respective allele. Validation of the targeted loci was performed on genomic DNA from F1 animals by RFLP, Sanger sequencing, and qPCR-based copy number assays. To exclude additional unwanted modifications, putative off-target sites of the Cd68-specific crRNA were predicted using the CRISPOR online tool.^68^ Predicted loci were PCR-amplified and verified by Sanger sequencing. Validated animals without any off-target mutations were used for further breeding. All animal experiments were approved by the Bavarian government under license number ROB-55.2Vet-2532.Vet_02-16-121. All mice were handled according to institutional guidelines approved by the animal welfare and use committee of the government of Upper Bavaria and housed in standard cages in a specific pathogen-free facility on a 12-h light/dark cycle with ad libitum access to food and water.

#### Isolation of murine adult primary microglia

Primary microglia were isolated using MACS Technology (Miltenyi Biotec) according to manufacturer’s instructions. Briefly, adult mice (C57BL/6j) were sacrificed by CO_2_ inhalation and brains collected in ice-cold DPBS with magnesium and calcium (Gibco). Brains were cut into smaller pieces and dissociated by enzymatic digestion using the Adult Brain Dissociation Kit (Miltenyi Biotec). Homogenates were filtered, debris removed, CD11b-positive microglia magnetically labeled with CD11b MicroBeads (Miltenyi Biotec), loaded onto MACS columns (Miltenyi Biotec) and subjected to magnetic separation. Cells were seeded on poly-L-lysine (Sigma) pre-coated culture dishes. After 1 day in culture, media was renewed. Cells were grown in culture for 7 days.

#### Transient transfection

For siRNA-mediated knockdown experiments, cells were reversely transfected with 30 nM of non-targeting or target-specific siRNA nucleotides (Dharmacon On-Target Plus) using Lipofectamine RNAiMAX (Invitrogen) according to manufacturer’s instructions. After 24 h, media was replaced and cells were kept in culture for a total of 72 h. To introduce plasmids, cells were transiently transfected when they reached at least 50 % confluency for 24 – 48 h using either polyethylenimine (PEI, Sigma) (stock: 1 mg/ml) or Lipofectamine 2000 in a 1:3 ratio (µg DNA: µl transfection reagent) according to manufacturer’s instructions.

#### Treatments

Cells were treated with L-Leucyl-L-Leucine methyl ester hydrobromide (LLOMe, Sigma) in a range from 250 µM up to 2 mM. Unless stated otherwise, 250 µM were used throughout this study. For recovery experiments, LLOMe was washed out twice with DPBS after 1 h of treatment and cells were kept in culture with fresh pre-warmed media for the indicated time. Other reagents used in this study are: BafA1 (Biomol, 200 nM, 2 h; 20 nM, 16 h), Bortezomib (LC Labs1 µM, 8 h), Monensin (Sigma-Aldrich, 50 µM, 2 h), DC661 (MedChem Express, 3 µM, 2 h), GPN (Cayman Chemical, 100 µM, 1 h), nanoSiO_2_ (Invivogen, 50 µg/ml, 6 h), Lysotracker Red DND-99 (Invitrogen, 100 nM).

#### Immunofluorescence staining and confocal microscopy

Cells were grown on glass coverslips. For BV2 cells and primary microglia, coverslips were coated with poly-L-lysine (Sigma). Cells were washed three times with DBPS (Gibco) and fixed with 4% paraformaldehyde (PFA, Santa Cruz) for 10 min at RT. Depending on the antibody, two different staining protocols were used. All steps were carried out at RT. For protocol 1, fixed cells were permeabilized with 0.5% Triton X-100 (Merck) in DPBS for 10 min and blocked with 1% BSA-DPBS for 1 h at RT. Primary and secondary antibodies were diluted in 0.1% BSA-DPBS blocking buffer and incubated successively in the dark for 1 h with DPBS washing steps in between. For protocol 2, permeabilization was performed with 0.1% Triton-X in DPBS for 10 min and blocking with blocking buffer (3% BSA, 0.1% Triton X-100, 0.1% Saponin (Sigma) in DPBS) for 1 h. Primary and secondary antibodies were used diluted in blocking buffer. For microglia, coverslips were treated for 1 min with TrueBlack (Biotium) (diluted 1:20 in 70% EtOH), washed and mounted with ProLong Gold Antifade with DAPI (Invitrogen). For Lysotracker assays, coverslips were mounted after fixation without permeabilization. Coverslips were imaged with an LSM 800 Carl Zeiss microscope using an oil-immersion objective at 63x magnification and ZENblue edition software. Images were processed with ZENblue 2011 (Zeiss, version 2.5).

#### Super-resolution microscopy using DNA-PAINT immunolabeling

GFP-RNF126 overexpressing cells were grown on poly-L-lysine coated chambered coverslips (Ibidi), treated with LLOMe for 1 h, and washed thrice with DPBS before fixation with 4% PFA at RT for 20 min. Samples were incubated in blocking buffer (1× DPBS, 0.05% NaN_3_, 2% BSA, 0.05 mg/ml sheared salmon sperm DNA) for 1 h at RT. Primary antibodies and the DNA-PAINT handle conjugated Anti-GFP nanobody (Nano-Tag) were added 1:200 into blocking buffer and incubated at 4°C overnight (ON). Samples were washed 3x 5 min with DPBS. DNA-PAINT handle conjugated secondary nanobody rabbit IgG (Nanotag) and secondary rat antibody (Jackson Immuno) were added 1:200 in blocking buffer and incubated for 90 min at RT. Conjugation of nanobodies to DNA-PAINT docking sites and labeling of anti-rat secondary antibody via a DBCO-sulfo-NHS ester linker was performed as previously described.^69^ Samples were washed 3x 5 min with DPBS, then post-fixed with 4% PFA for 7 min at RT. Samples were then washed with DPBS and quenched with 0.2 M NH_4_CL for 5 min at RT. A 1:4 dilution of Gold nanoparticles (90 nm, Cytodiagnostics) was incubated for 5 min at RT. For DNA-PAINT Imaging: DNA-PAINT imager strands were diluted to the appropriate imaging concentration in Buffer C (500 mM NaCl in 1× DPBS pH 7.4) containing the oxygen scavenger system PCA/PCD/Trolox and added to the sample for sequential DNA-PAINT imaging, i.e. Exchange-PAINT.^70^ Bright field and GFP imaging were taken before DNA-PAINT data acquisition to help find the correct focal plane and access cell viability. HILO illumination was used for data acquisition. Fluorescence imaging for DNA-PAINT experiments was carried out on an inverted microscope (Nikon Instruments, Eclipse Ti2) with the Perfect Focus System, applying an objective-type TIRF configuration equipped with an oil-immersion objective (Nikon Instruments, Apo SR TIRF ×100, numerical aperture 1.49, oil). 488 nm and 560 nm lasers (MPB Communications, 1 W) were used for excitation and coupled into the microscope via a Nikon manual TIRF module. The laser beams were passed through cleanup filters (Chroma Technology, ZET488/10× for 488 nm excitation, ZET561/10× for 560 nm excitation, and coupled into the microscope objective using a beam splitter (Chroma Technology, ZT488rdc-UF2 for 488 nm excitation, ZT561rdc-UF2 for 560 nm excitation). Fluorescence was spectrally filtered with an emission filter (Chroma Technology, ET525/50 m and ET500lp for 488 nm excitation, ET600/50 m and ET575lp for 560 nm) and imaged on an sCMOS camera (Hamamatsu, ORCA-Fusion BT) without further magnification, resulting in an effective pixel size of 130 nm (after 2 × 2 binning). The central 1,152 × 1,152 pixels (576 × 576 after binning) of the camera were used as the region of interest, and the scan mode was set to ‘ultra quiet scan’. Raw microscopy data were acquired using μManager^38^ (Version 2.0.1). Raw fluorescence data from DNA-PAINT imaging was subjected to super-resolution reconstruction using the “Picasso” software package (latest version available on https://github.com/jungmannlab/picasso).^71^ Drift correction was performed with a redundant cross-correlation and gold nanoparticles as fiducials. Gold nanoparticles were also used for alignment of multi-target Exchange-PAINT images.^70^

#### Live cell microscopy and photo damage

Live cell imaging was performed at 37 °C in imaging medium (PAN-Biotech) supplemented with 10% FCS and 1% Pen/Strep. Spinning disk microscopy with laser induced lysosomal damage was performed on the Eclipse Ti-E (Nikon) inverted microscope equipped with an Andor AOTF Laser Combiner, a CSU-X1 Yokogawa spinning disk unit and an iXon3 897 single photon detection EMCCD camera (Andor Technology). Laser lines used for excitation of EGFP and mCherry were 488 nm and 561 nm, respectively. Images were acquired using a CFI APO TIRF 100x/1.49NA oil immersion objectives (Nikon). To induce damage, cells were loaded with 125 nM AlPcS2a (Frontier Scientific) as described before.^21^ Defined regions in cells were exposed to a 640 nm laser with 100% power for 4 x 1000 µs pixel dwelling time using Andor Revolution FRAP and Photo Activation illumination system (FRAPPA). Image acquisition was controlled by Andor IQ3 Software (Andor Technology) as previously described.^21^

#### Transmission electron microscopy

Parental and RNF126 KO cells were grown on plasma-treated ACLAR® plastic films (Science Services). After mock or LLMOe treatment, cells were washed twice in DPBS and media was replaced. Subsequently, prewarmed 2x fixative (5% glutaraldehyde (EM-grade, Science Services), 0.2 M cacodylate buffer pH 7.4 (Science Services)) was added 1:1 to the culture medium. After 5 min this mixture was replaced by 1x fixative (2.5% glutaraldehyde in 0.1 M cacodylate buffer) and incubated for 25 min on ice. Postfixation and -contrasting was performed by several washes in 0.1 M cacodylate buffer on ice followed by incubation in 1% osmium tetroxide (Science Services) and 0.8% potassium ferrocyanide (Sigma Aldrich) in 0.1 M sodium cacodylate buffer. Cells were further contrasted using 0.5% uranyl acetate (Science Services) in water. After dehydration in ethanol and infiltration in LX112 resin (Ladd Research) the cell monolayer was embedded into resin blocks and cured for 48 h at 60 °C. Ultrathin sections (80 nm thick) were generated on a Leica UC6 ultramicrotome and adhered to formvar-coated copper grids (Plano). The sections were imaged on a JEM 1400plus (JEOL) transmission electron microscope using a XF416 camera (TVIPS) and the EM-Menu software (TVIPS). Image analysis was conducted using Fiji. Experiments were performed in biological triplicates. For quantification analysis multiple t-test with False Discovery Rate (FDR) approach (two-stage step-up method of Benjamini, Krieger and Yekutieli; FDR<0.05) was used.

#### Electron tomography

Grids were imaged on a Tecnai F30 (Thermo Fisher Scientific, Waltham, MA, USA) transmission electron microscope equipped with a Gatan OneView camera (Gatan/Ametek Pleasanton, CA, USA). Precise target positions were manually selected using the SerialEM software and acquired by single-axis tomography (−60° to +60° per axis; increment of 1°) with a pixel size of 1.56 nm/px.^72^ Tilt series were reconstructed using the Etomo from the IMOD software package.^73^ Image segmentation was performed in VAST and the models were rendered using Blender.^74^

#### Flow cytometry

Analysis of lysophagy flux in parental and RNF126 KO cells expressing mKeima-Gal3 was performed as described previously.^75^ In brief, cells were treated with 1 mM LLOMe for 1 h followed by different recovery time points. As negative control 20 nM Bafilomycin was added to block lysophagy. Cells were harvested after indicated time points by trypsinization and live singlets analyzed for ratio of mean fluorescence intensity (MFI) of acidic Keima signal (mCherry) to neutral Keima signal (PercP) with a BD FACS Melody flow cytometer. Experiments were performed in biological triplicates. For statistical analysis multiple t-test with False Discovery Rate (FDR) approach (two-stage step-up method of Benjamini, Krieger and Yekutieli; FDR<0.05) was used.

#### Cell lysis and Immunoblotting

Cell pellets were lysed in RIPA buffer (50 mM Tris-HCl pH 7.5, 150 mM NaCl, 0.1% SDS, 0.5% sodium deoxycholate, 1% NP-40 / 1% Triton X-100, 1x protease inhibitor (Roche) and 1x PhosStop (Roche)) on ice for 30 min. Lysates were cleared via centrifugation (21,130 x g 4 °C, 10 min). Protein concentration was determined using BCA and adjusted between samples if no prior cell count was performed. Lysates were boiled in 3x sample loading buffer (200 mM Tris-HCl pH 6.8, 6% SDS, 20% glycerol, 0.1 g/ml DTT and 0.01 mg/ml Bromophenol Blue) at 95 °C for 5 min. Proteins were size-separated by SDS-PAGE and transferred onto 0.45 µm nitrocellulose (Cytiva, Amersham Protran) or methanol-activated 0.45 µm PVDF membranes (Merck). Immunoblots were blocked with either 5% low-fat milk (Roth), 3% BSA (Sigma) or 0.2% I-Block (Invitrogen) in TBS supplemented with 0.1% Tween-20 (TBS-T) according to antibody requirements at RT for 1 h. Membranes were incubated with primary antibodies diluted in blocking buffer at 4 °C ON shaking, followed by several wash steps with TBS-T and incubation with secondary HRP-conjugated antibody at RT for 1 h. Secondary antibodies were diluted in either 0.5% milk-TBS-T, 0.3% BSA-TBS-T or 0.2% I-Block-TBS-T. After repeated washing, immunoblots were analyzed by enhanced chemiluminescence using Western Lightning Plus ECL (Perkin Elmer) and X-ray films (Fuji Medical).

#### HA immunoprecipitation

Cells were either lysed in mammalian cell lysis buffer (MCLB) (50 mM Tris-HCl pH 7.5, 150 mM NaCl, 0.5% NP-40, 1x protease inhibitor (Roche) and 1x PhosStop (Roche)) or Glycerol buffer (20 mM Tris-HCl pH 7.5, 150 mM NaCl, 0.5% Triton X-100, 5 mM EDTA pH 8.0, 10% glycerol, 1× protease inhibitor (Roche) and 1x PhosStop (Roche)) on ice for 30 min followed by clearing and protein concentration adjustment. An aliquot of the cleared lysate was saved as input before incubating with pre-equilibrated anti-HA agarose beads (Pierce) overhead-rotating at 4 °C ON. The input was boiled with 3x sample loading buffer at 95 °C for 5 min. Next day, IP samples were washed five times with ice-cold MCLB or respective Glycerol buffer. Beads were dried and boiled in 3x sample loading buffer at 95 °C for 5 min. Input and IP samples were analyzed by immunoblotting.

#### Denaturing GFP immunoprecipitation

GFP chimera expressing parental and RNF26 KO cells were treated with 1 mM LLOMe for 1 h and chased for additional 2 h. To detect ubiquitylation of GFP-chimeras, cells were denatured in TSD buffer (50 mM Tris-HCl, pH 7.5, 1% SDS, 5 mM DTT) and then diluted with TNN buffer (20 mM Tris-HCl, pH 7.5, 200 mM NaCl, 0.5% NP-40, 1× protease inhibitor (Roche).^76^ The cleared lysates were incubated with GFP-coupled Sepharose beads to pull down GFP-tagged proteins (1 h at 4 °C). Beads were washed 3x with ice-cold TNN buffer containing 0.1% SDS and further processed by SDS PAGE and Western blotting.

#### APEX2-mediated biotinylation

APEX2-chimera expressing cells were treated with 500 µM biotin-phenol (BP) (IrisBiotech) at 37 °C for 30 min prior to a 1 min pulse with 1 mM H_2_O_2_ (Sigma) at RT. Peroxidase activity was immediately quenched by decanting the media and washing the cells three times with ice-cold quencher solution (10 mM sodium azide, 10 mM sodium ascorbate and 5 mM Trolox (Sigma) in DPBS.^28^ Afterwards cells were washed three times with ice-cold DPBS following harvest with 0.25% Trypsin-EDTA (Gibco). Protein amount was either equalized between samples by adjusting cell counts (for MS experiments) before lysis or via BCA after RIPA lysis (for immunoblotting).

#### Streptavidin pulldown

Biotin-phenol treated APEX2-chimera expressing cells were lysed in quenching-RIPA buffer (50 mM Tris-HCl pH 7.5, 150 mM NaCl, 0.1% SDS, 0.5% sodium deoxycholate, 1% Triton X-100, 1x protease inhibitor (Roche), 1x PhosStop (Roche), 10 mM sodium azide, 10 mM sodium ascorbate (Sigma), 5 mM Trolox) on ice for 30 min. Lysates were cleared by centrifugation and supernatants were incubated ON with pre-equilibrated streptavidin agarose beads (Sigma) overhead-rotating at 4 °C. Beads were subsequently washed twice with RIPA buffer, four times with freshly prepared 3 M Urea buffer (3 M Urea, 50 mM ABC (Sigma) and suspended in defined volume of 3 M Urea buffer. Samples were reduced with 5 mM TCEP (Sigma) at 55 °C for 30 min shaking, alkylated with 10 mM IAA (Sigma) at RT for 20 min and quenched with 20 mM DTT (Sigma) at RT. Following two washes with freshly prepared 2 M Urea buffer (2 M Urea, 50 mM ABC), beads were suspended in 2 M Urea buffer and proteins digested with 1 μg MS-grade trypsin (Promega) at 37 °C ON. Peptides were collected by pooling the supernatant with two additional 2 M Urea buffer washes, immediately acidified with 1% TFA (Fluka) and concentrated by vacuum centrifugation.^77^ Digested peptides were desalted on custom-made C18 stage tips and solved in buffer A (0.1% FA (Merck) in MS-grade water (Roth)).^78^

#### Lysosome immunoprecipitation

TMEM192-3xHA expressing cells were processed essentially as described before.^35^ Cells were washed twice in ice-cold DPBS and harvested via scraping in DBPS supplemented with protease inhibitors. Cell count was adjusted and cells suspended in ice-cold Lyso-IP buffer (50 mM KCl, 100 mM KH_2_PO_4_, 100 mM K_2_HPO_4_ pH 7.2, protease inhibitors). Following dounce homogenization with a tight-fitting pestle, homogenates were cleared from unbroken cells and nuclei via centrifugation (1,500 x g, 10 min, 4 °C) and supernatant was incubated with pre-equilibrated magnetic anti-HA beads (Thermo Scientific) on an overhead-rotator at 4 °C for 1 h. Beads were subsequently washed three times in Lyso IP buffer supplemented with 300 mM NaCl and lysosomes eluted by addition of freshly prepared 8 M Urea buffer (8 M Urea, 50 mM Tris-HCl pH 8.0, 150 mM NaCl) shaking at 4 °C for 30 min. The eluate was then sonicated and sequentially treated with 5 mM DTT shaking at 55 °C for 30 min, 15 mM IAA for 30 min at RT, and 10 mM DTT for 15 min at RT. Proteins were extracted with methanol-chloroform precipitation by sequentially mixing the sample with 600 μl methanol (Roth), 150 μl chloroform (Merck) and 450 μl MS-grade water, centrifugation (14,000 x g, 5 min, RT), removing the hydrophilic phase, washing with 450 µl methanol and finally pelleting the proteins via centrifugation (14,000 x g, 5 min, RT). The protein pellet was dried by vacuum centrifugation before suspending it in 50 mM ABC buffer and ON digestion with 1 µg trypsin at 37 °C. Digests were stopped with 5 % ACN (Roth)/ 5% FA. Peptides were dried by vacuum centrifugation, reconstituted in buffer C (5% ACN, 1% TFA), desalted on custom-made C18 stage tips and solved in buffer A.

#### diGly proteomics

Two conditions were compared by SILAC. Cells were processed essentially as described before.^79^ Cells were washed twice with ice-cold DPBS and immediately scraped in 5 ml denaturing lysis buffer (9 M urea, 50 mM Tris-HCl pH 8.0, 150 mM NaCl, 50 µM DUB inhibitor PR-619 (Millipore), protease inhibitors). After multiple rounds of sonication on ice, non-solubilized material was removed via centrifugation (15,000 x g, 10 min, 15 °C) and protein concentration determined with BCA. Heavy-and light-labeled protein lysate was combined in a 1:1 ratio to obtain a total amount of 10 mg protein per sample. Proteins were then prepared for digest by sequentially adding reducing agent (5 mM DTT, 1 h, 22 °C, 200 rpm), alkylating agent (10 mM chloroacetamide, 1 h, 22 °C, 200 rpm) and quencher (5 mM DTT). Afterwards, samples were diluted 1:1 with 50 mM Tris-HCl (pH 8.0) and proteins digested with 5 ng/µl Lys-C (Wako) for 1 h at 22 °C. Subsequently, samples were diluted 1:3 with 50 mM Tris-HCl (pH 8.0), 1 mM CaCl_2_ was added and samples digested with 50 µg trypsin ON at 37 °C. Digests was stopped by acidification (0.4 % TFA). Peptide mixture was cleared from insoluble material with centrifugation (2,500 x g, 10 min, RT) and soluble fraction concentrated and desalted with reverse-phase C18 Sep-Pak extraction cartridges (Waters) followed by lyophilization as described previously.^80^ Dried peptides were reconstituted in immunoaffinity purification (IAP) buffer (50 mM MOPS (pH 7.4), 10 mM Na_2_HPO_4_, 50 mM NaCl) and centrifuged (2,500 xg, 5 min, 4 °C). 0.1 % of peptide mixture was separated as input fraction and residual supernatant was incubated with pre-equilibrated anti-ubiquitin remnant motif (K-ε-GG) agarose beads (Cell Signaling) at 4 °C overhead-rotating for 1 h. For maximal capture, each sample was immunoprecipitated sequentially two times and IP fractions were processed individually. Beads were washed thrice with ice-cold IAP buffer and once with ice-cold DBPS before elution of bound peptides with 0.15 % TFA using Micro Bio-Spin^TM^ columns (Biorad). Elution was performed four times sequentially. Input and IP fractions were desalted on custom-made C18 stage tips and solved in buffer A before applying it to LC-MS/MS.

#### MS data collection and analysis

All samples were separated using an Easy-nLC1200 liquid chromatograph (Thermo Scientific) followed by peptide detection on a Q Exactive HF mass spectrometer (Thermo Scientific). Peptides were loaded onto 75 μm × 15 cm custom-made fused silica capillaries packed with C18AQ resin (Reprosil-PUR 120, 1.9 μm, Dr. Maisch HPLC). Flow rates and gradients were adjusted to the respective experiment accordingly. For Lyso-IP and streptavidin pulldown samples, peptide mixtures were separated on a 35 min buffer B (80 % ACN, 0.1 % FA) gradient (10–38% for 23 min, 38–60% for 2 min, 60–100% for 2 min, 100% for 3 min, 100%–5% for 2 min, 5% for 3 min) at a flow rate of 400 nl/min. For whole cell proteomics (input of diGly samples), peptides were separated on a 140 min buffer B gradient (3–6% for 2 min, 6-30% for 90 min, 30–44% for 20 min, 44–75% for 10 min, 75–100% for 5 min, 100% for 5 min, 100–3% for 5 min, 3% for 3 min) at a flow rate of 250 nl/min. For diGly IP samples, peptides were separated on a 60 min buffer B gradient (10–38% for 35 min, 38-60% for 5 min, 60–100% for 5 min, 100% for 5 min, 100-5% for 5 min, 5% for 5 min) at a flow rate of 400 nl/min. Peptides were ionized using a Nanospray Flex Ion Source (Thermo Scientific) and identified in fullMS / ddMS² mode. For Lyso-IP and streptavidin pulldown experiments, the top 15 most intense peaks from each full MS scan were selected for subsequent fragmentation, with a dynamic exclusion of 20.0 s excluding peptides with unassigned charge or charges of 1 or >8. MS1 resolution was set to 60,000 with a scan range of 300–1650 m/z, MS2 to 15,000. AGC target1 was set to 3e6, AGC target2 to 1e5. For whole cell proteomics (input diGly samples), the top 20 most intense peaks from each full MS scan were selected for subsequent fragmentation, with a dynamic exclusion of 120.0 s excluding peptides with unassigned charge or charges of 1 or >8. For whole proteome analysis, MS1 resolution was set to 120,000 with a scan range of 300–1700 m/z, MS2 to 15,000. AGC target1 was set to 3e6, AGC target2 to 1e5. For diGly IP samples, the top 5 most intense peaks were picked for fragmentation, with a dynamic exclusion of 40.0 s excluding peptides with unassigned charge or charges of 1, 2 or >8. MS1 resolution was set to 60,000 with a scan range of 300–1750 m/z, MS2 to 30,000. AGC target1 was set to 1e6, AGC target2 to 5e5. Data collection was controlled by Tune/Xcalibur (Thermo Scientific). Raw data files were analyzed using MaxQuant (1.6.0.1) Andromeda search engine in reversed decoymode based on a human reference proteome (Uniprot-FASTA, UP000005640, downloaded September 2017) with an FDR of 0.01 on the protein level.^81, 82^ Digestion parameters were set to specific digestion with trypsin/p with a maximum number of 2 missed cleavage sites and minimum peptide length of 7. For all experiments, methionine oxidation (15.994946) and N-terminal protein acetylation (42.010565) were set as variable modifications and carbamidomethylation of cysteine as fixed modification. For diGly MS experiments, diGly remnant (114.042927; excluded from the C terminus) was set as additional variable modification. The peptide mass tolerance was set to 20 ppm (first search) and to 4.5 ppm (main search). For label-free experiments, label-free quantification (with a minimum ratio count set to 2) was selected, whereas for SILAC experiments, multiplicity was set to 2 with light Lys0 and heavy Lys8 labeling. Re-quantification and match-between runs were selected for all analysis. Output text files from MaxQuant analysis were further processed in Perseus (version 1.6.14.0).^83^ For whole cell proteomics, Lyso-IP and APEX2 samples, protein groups files were used for further analysis. For ubiquitin remnant profiling the analysis was performed on the diGly modified peptides file. Briefly, common contaminants, site-specific and reverse identifications were excluded. Protein groups with less than 2 peptide and MS/MS counts were removed. Furthermore, data was filtered to keep only protein groups / peptides present in at least three out of four replicates per condition. LFQ intensities and SILAC ratios were log_2_ transformed. FOR LFQ experiments, remaining missing values were imputed with random numbers from a normal distribution with 0.3 width and 1.8 down shift calculated separately for each sample. For SILAC experiments, no imputation was performed. Statistical analysis of Lyso-IP and APEX2 experiments were performed with a two-sided two-sample Student’s t-test with permutation-based FDR (q-value<0.05). For SILAC diGly-MS experiments, one-sample t-test (p-value<0.05) with multiple comparison correction (Benjamini-Hochberg correction FDR<0.05) (q-value<0.05) was done. DAVID 2021 (https://david.ncifcrf.gov/) was used for functional enrichment analysis.^84, 85^

---

**Supplementary Figure S1.**
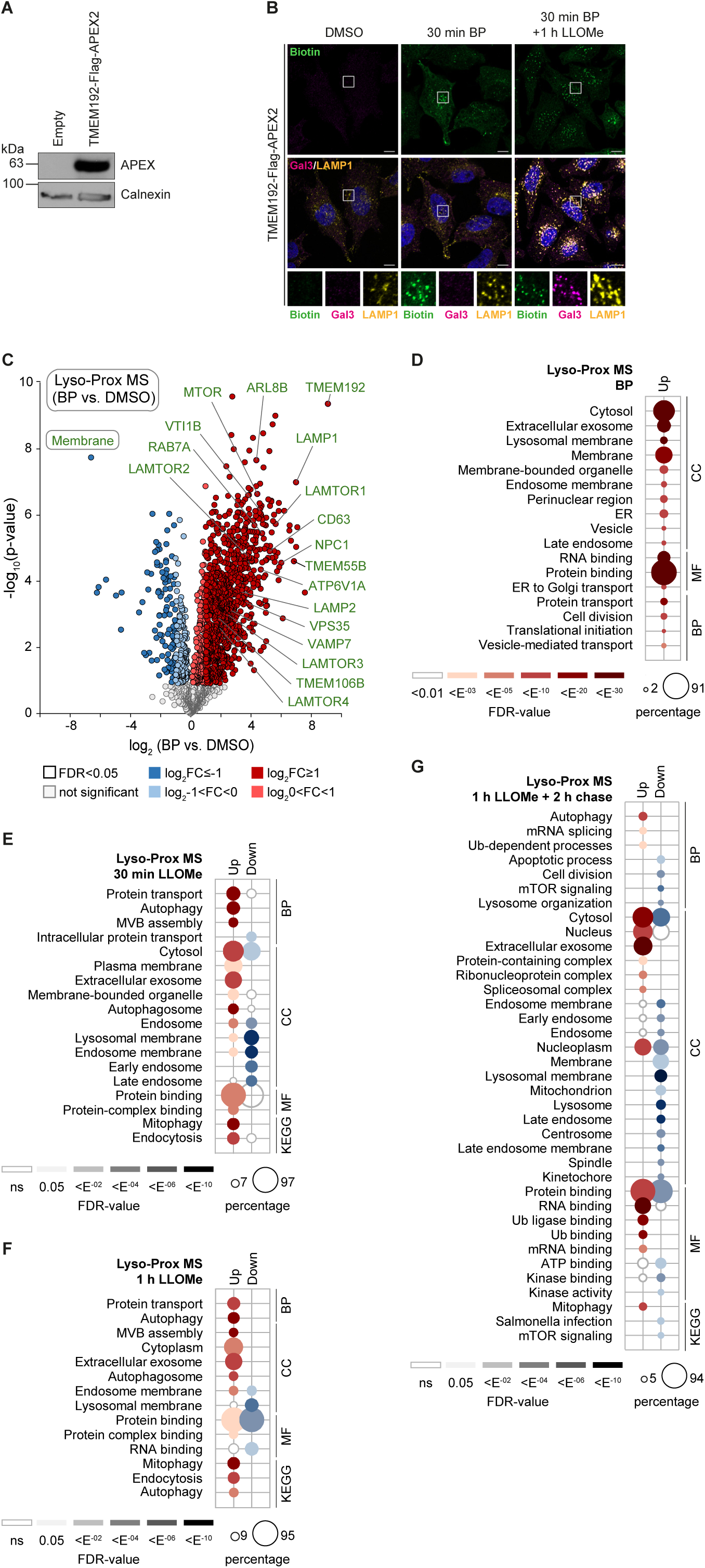
Proximity labeling of damaged lysosomes. (A) Immunoblot of lysates from empty or TMEM192-Flag-APEX2 expressing cells. (B) TMEM192-Flag-APEX2 cells treated as indicated were fixed and stained for indicated antibodies. Insets show magnification of boxed areas. Scale bars, 10 µm. (C) Volcano plot of proteins enriched upon biotinylation induction. Cells were treated with BP or DMSO. Experiments were performed in biological quadruplicates and statistical analysis using two-sided two sample Student’s t-test (p-value<0.05) with permutations-based correction (FDR<0.05). Significantly altered proteins (FDR<0.05) are labeled in dark (log_2_FC ≥ I1I) and light color (log_2_I0<FC<1I), respectively. Lysosomal membrane-associated proteins are highlighted in green. (D) GO enrichment analysis of proteins significantly enriched at lysosomes (FDR<0.05, log_2_FC>0) from (C). Dot size correlates to number of proteins, dot color to term enrichment (FDR). BP, biological process; CC, cellular component, MF, molecular function; KEGG, Kyoto Encyclopedia of Genes and Genomes. (E-G) GO enrichment analysis of proteins significantly enriched or diminished (FDR<0.05, log_2_FC ≥ I1I) upon 30 min LLOMe treatment (E), 1 h LLOMe treatment (F) or 1 h LLOMe treatment followed by 2 h chase (G).

**Supplementary Figure S2.**
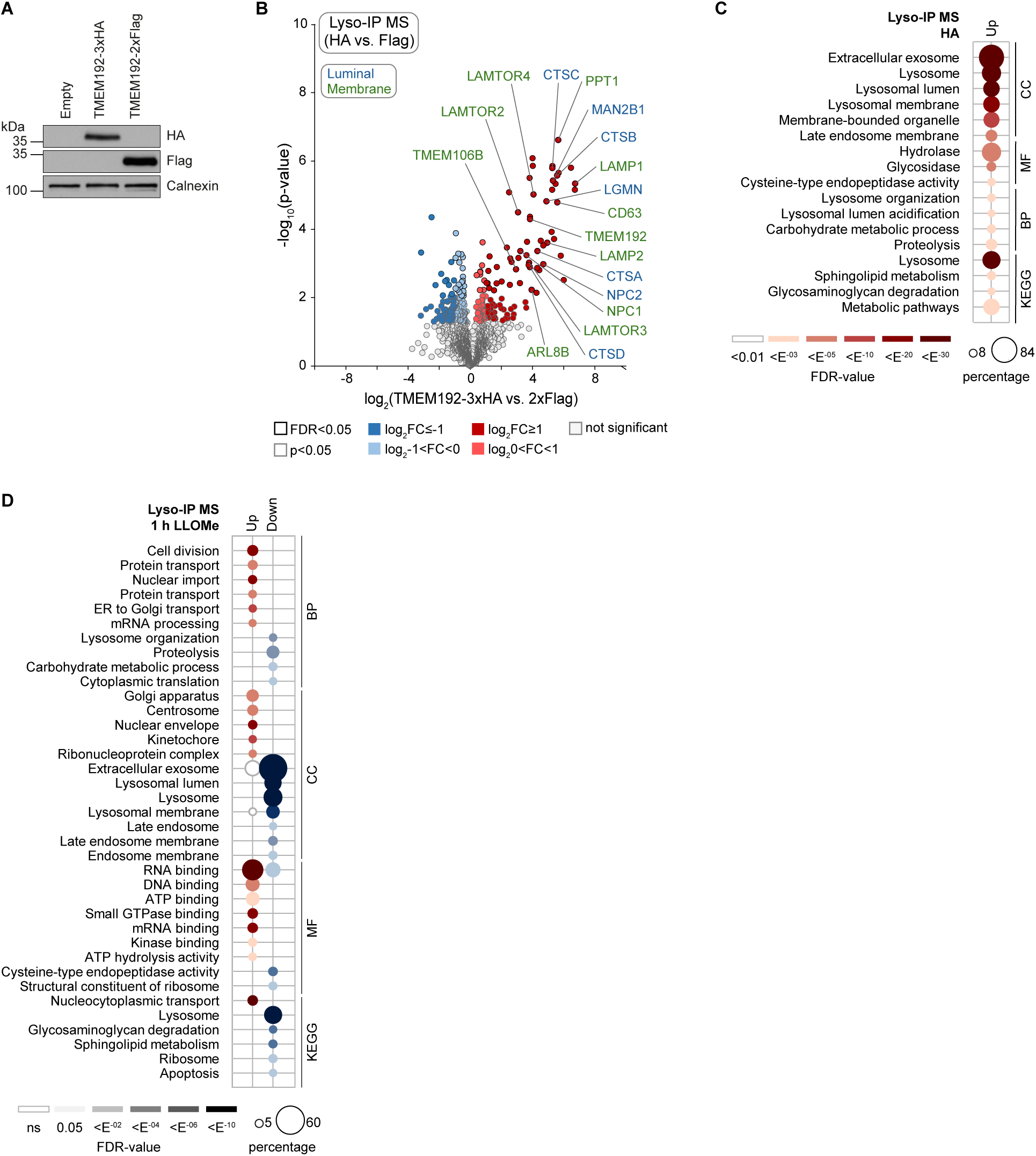
Immune isolation of damaged lysosomes. (A) Immunoblot of lysates from empty, TMEM192-3xHA or TMEM192-2xFlag expressing cells. (B) Volcano plot of proteins enriched at lysosomes. Homogenates from TMEM192-3xHA or TMEM192-2xFlag (negative control) cells were subjected to immune-isolation with anti-HA antibody. Experiments were performed in biological quadruplicates and statistical analysis using two-sided two sample Student’s t-test (p-value<0.05) with permutations-based correction (FDR<0.05). Significantly altered proteins (FDR<0.05) are labeled in dark (log_2_FC ≥ I1I) and light color (log_2_I0<FC<1I), respectively. Lysosomal membrane and luminal proteins are highlighted in green and blue, respectively. (C) GO enrichment analysis of enriched proteins (FDR<0.05 and log_2_FC>0) from (B). Dot size correlates to number of proteins, dot color to term enrichment (FDR). BP, biological process; CC, cellular component, MF, molecular function; KEGG, Kyoto Encyclopedia of Genes and Genomes. (D) GO enrichment analysis of lysosomal proteins significantly altered upon LLOMe (FDR<0.05 and log_2_FC ≥ I1I) from Figure 1D.

**Supplementary Figure S3.**
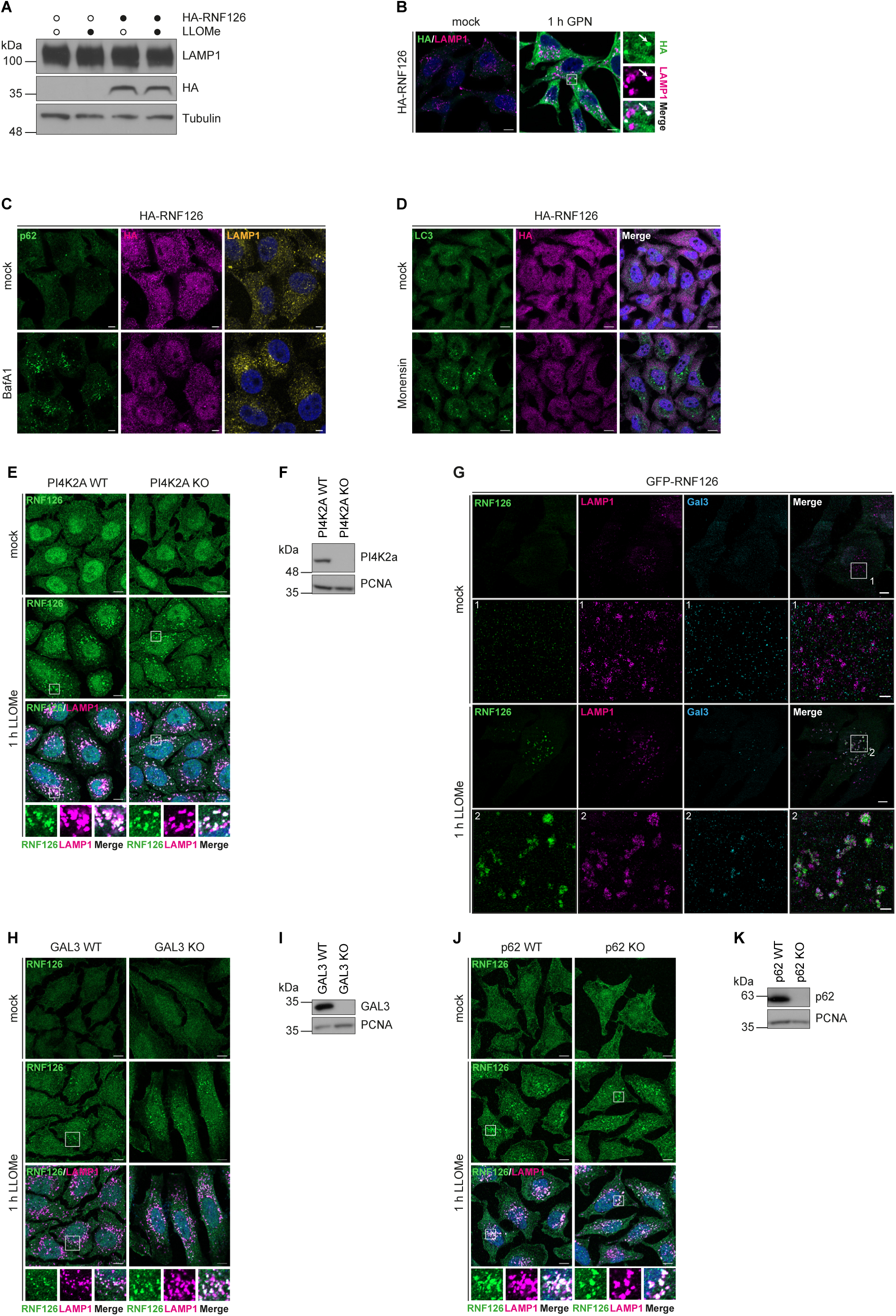
Requirements of RNF126 translocations to lysosomes. (A) Empty and HA-RNF126 expressing cells were treated with LLOMe, lysed and analyzed by immunoblotting with indicated antibodies. (B) HA-RNF126 cells were treated with GPN (100 µM), fixed and stained with anti-HA and -LAMP1 antibodies. Insets show magnification of boxed areas. Scale bars, 10 µm. (C) HA-RNF126 cells were treated with BafA1 (200 µM) for 2 h, fixed and stained with anti-HA and p62 antibodies. Scale bars, 5 µm. (D) HA-RNF126 cells were treated with Monensin (50 µM) for 2 h, fixed and stained with anti-HA and -LC3 antibodies. Scale bars, 10 µm. (E) PI4K2A WT and KO HeLa were treated with LLOMe, fixed and stained with indicated antibodies. Scale bars, 10 µm. (F) Immunoblot of lysates from PI4K2A WT and KO cells. (G) Cells expressing GFP-RNF126 were treated with LLOMe, fixed, immunolabeled with anti-LAMP1 and -Gal3 antibodies and subjected to DNA-PAINT technology for super-resolution microscopy. Scale bars, 5 µm. Insets show magnification of boxed areas. Scale bars of insets, 1 µm. (H) GAL3 WT and KO cells were treated with LLOMe, fixed and stained with indicated antibodies. Scale bars, 10 µm. (I) Immunoblot of lysates from GAL3 WT and KO cells. (J) p62 WT and KO cells were treated with LLOMe, fixed and stained with indicated antibodies. Scale bars, 10 µm. (K) Immunoblot of lysates from p62 WT and KO cells.

**Supplementary Figure S4.**
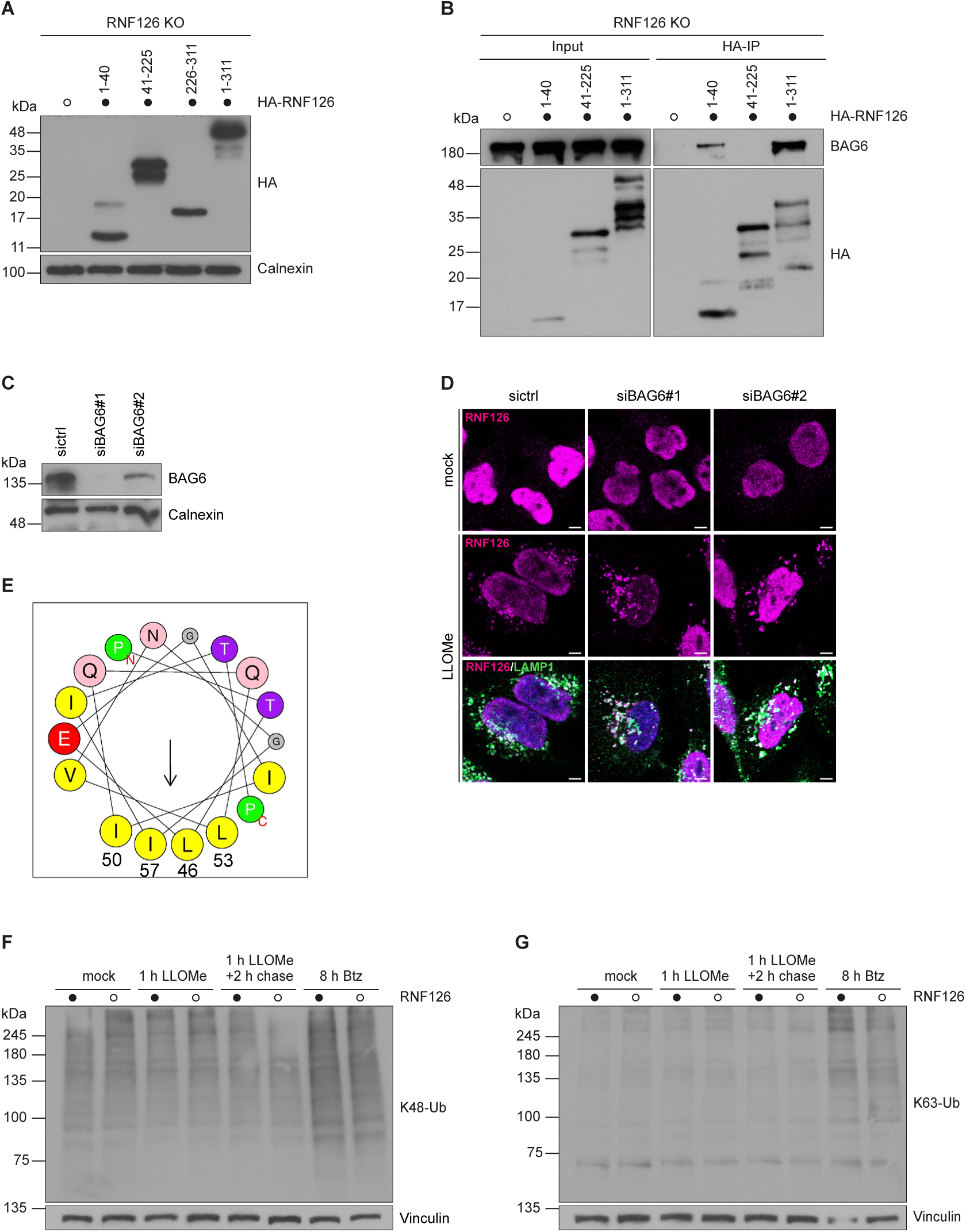
BAG6 is not required for lysosomal translocation of RNF126. (A) Reconstitution of indicated HA-RNF126 variants in RNF126 KO HeLa cells. (B) Lysates of RNF126 KO cells expressing indicated HA-RNF126 variants were subjected to HA-IP and immunoblotting with anti-HA and -BAG6 antibodies. (C) Cells were reversely transfected with BAG6 or non-targeting (sictrl) siRNAs (30 nM siRNA, 72 h) followed by lysis and immunoblotting. (D) BAG6-depleted cells (from C) were treated with LLOMe, fixed and stained with anti-RNF126 and -LAMP1 antibodies. Scale bars, 5 µm. (E) Model of amphipathic helix comprising amino acids leucine 46, isoleucine 50, leucine 53 and isoleucine 57 (LILI motif) potentially formed in presence of lipids. (F and G) Lysates from parental and RNF126 KO cells treated as indicated were analyzed by immunoblotting with anti-K48 and -K63 Ub antibodies.

**Supplementary Figure S5.**
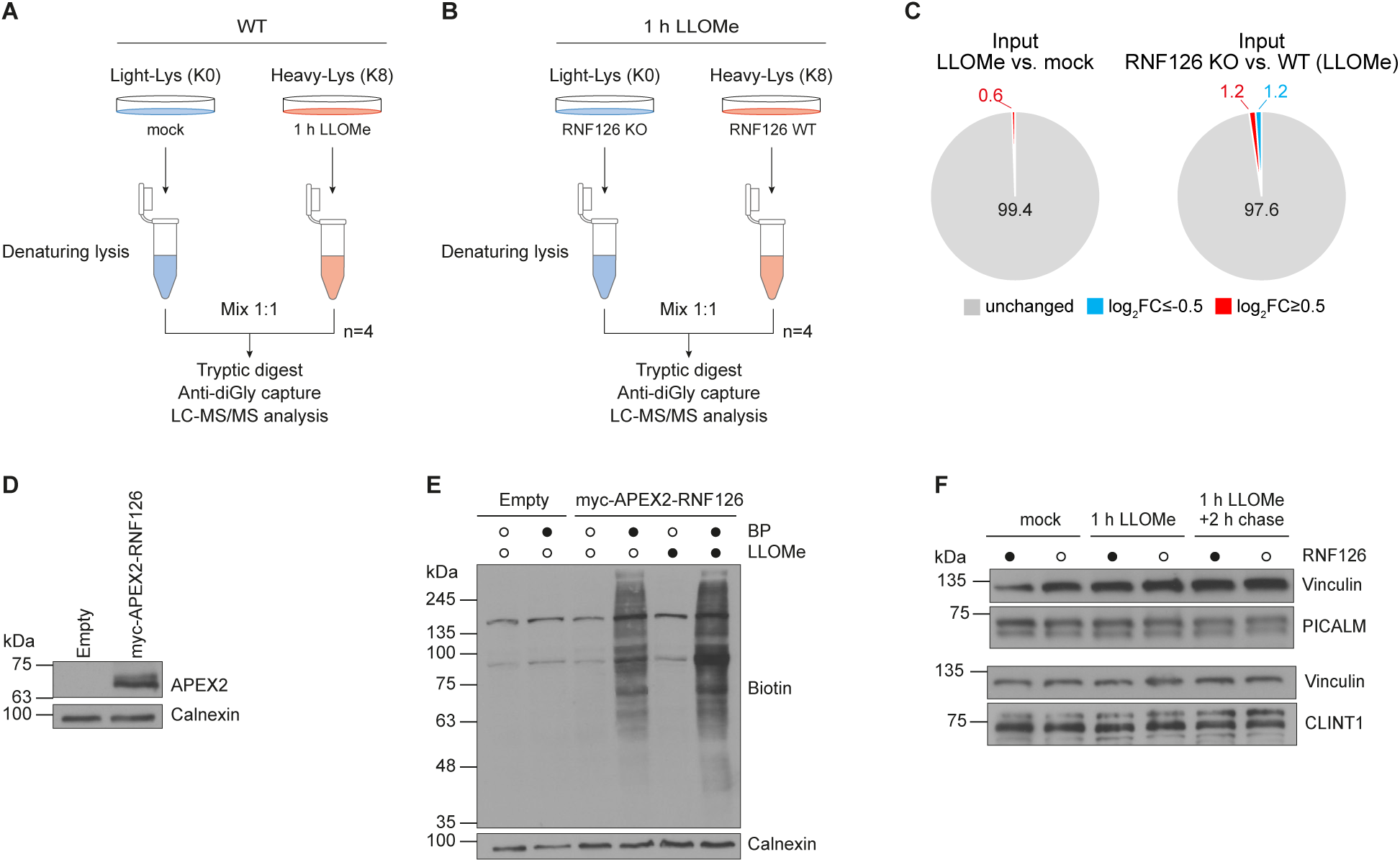
RNF126 substrate identification. (A and B) Experimental workflow of SILAC-based diGly MS to identify damage-induced ubiquitylation events (A) or RNF126-dependent ubiquitylation substrates under damage condition (B). Experiments were performed in biological quadruplicates and statistical analysis using one sample Student’s t-test (p-value<0.05) with multiple comparison correction (Benjamini-Hochberg correction FDR<0.05). (C) Percentage of quantified proteins whose abundance changed by at least twofold (log_2_ SILAC ratio >1 or <-1). (D) Immunoblot of lysates from empty or myc-APEX2-RNF126 expressing cells. (E) Myc-APEX2-RNF126 cells were incubated 30 min with BP and pulsed with H_2_O_2_ in mock-and LLOMe-treated condition. Cells were lysed and analyzed for biotinylation. (F) RNF126 WT and KO cells were treated as indicated, lysed and analyzed by immunoblotting.

**Supplementary Figure S6.**
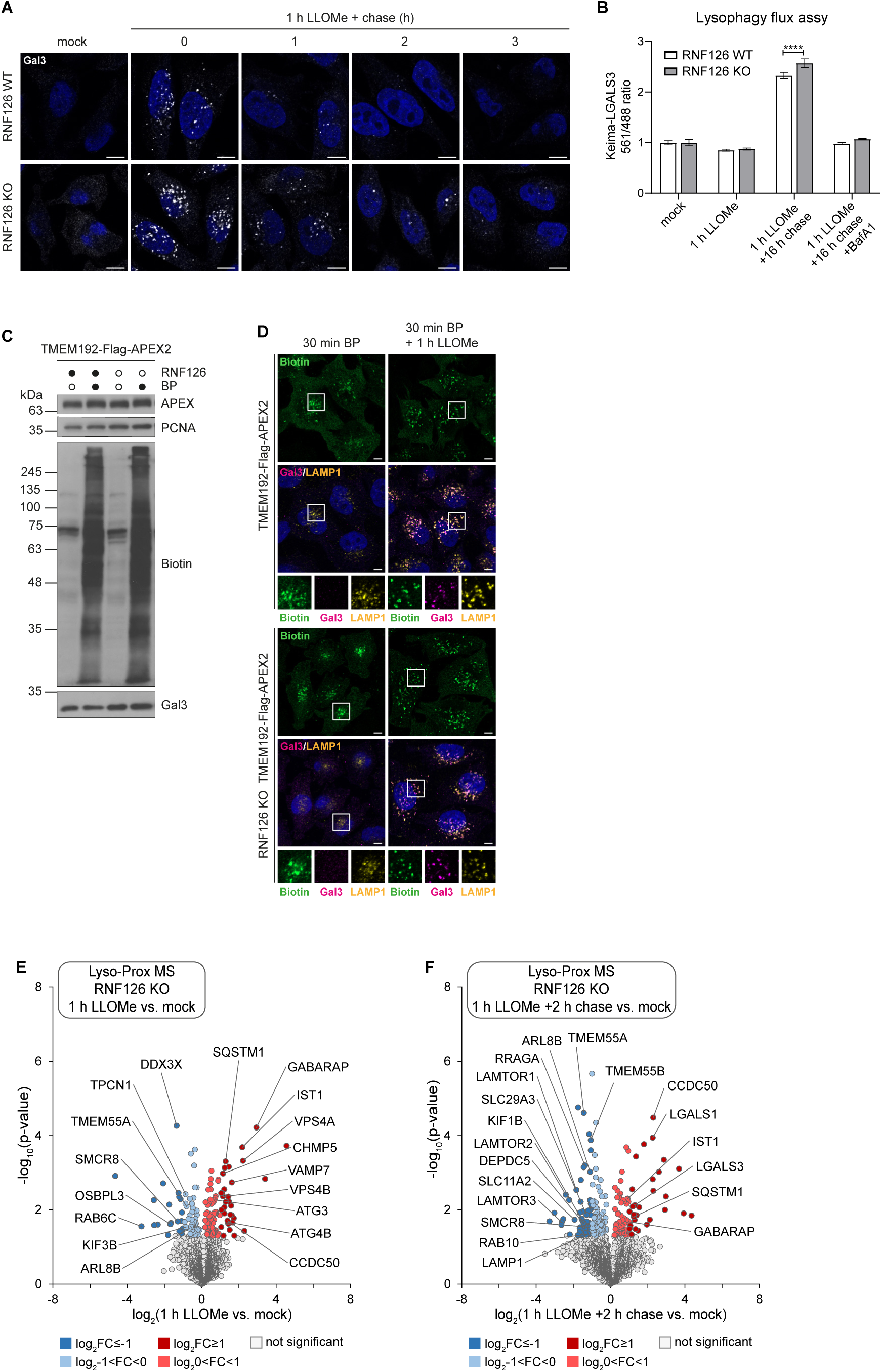
Lysosomal proximity proteomics in RNF126 KO cells. (A) RNF126 WT and KO cells were treated with LLOMe followed by a washout and 4 h chase with fresh media. Cells were fixed and stained with an anti-Gal3 antibody. Scale bars, 5 µm. (B) Lysophagy flux assay. Triplicate RNF126 KO and WT cells expressing Keima-LGALS3 were either left untreated or treated with LLOMe for 1 h followed by washout (16 h). Cells were subjected to flow cytometry to measure the 561 nm/488 nm ratio. All values are normalized to the untreated sample. ****p < 0.0001. The plot represents mean and standard deviation from three biological replicates. BafA1 (20 nM) was added post LLOMe treatment during washout as negative control. (C and D) RNF126 WT and KO cells expressing TMEM192-Flag-APEX2 were treated 30 min with BP and pulsed with H_2_O_2_. Cells were lysed and analyzed by immunoblotting (C) or fixed and stained with indicated antibodies (D). Scale bars, 10 µm. (E and F) Volcano plots of lysosomal proximity proteome changes in RNF126 KO cells upon damage. RNF126 KO cells were treated with LLOMe for 1 h (E) or for 1 h followed by washout and 2 h chase with fresh media (F). Experiments were performed in biological quadruplicates and statistical analysis using two-sided two sample Student’s t-test (p-value<0.05) with permutations-based correction (FDR<0.05). Significantly altered proteins (p<0.05) are labeled in dark (log_2_FC ≥ I1I) and light color (log_2_I0<FC<1I), respectively.

**Supplementary Figure S7.**
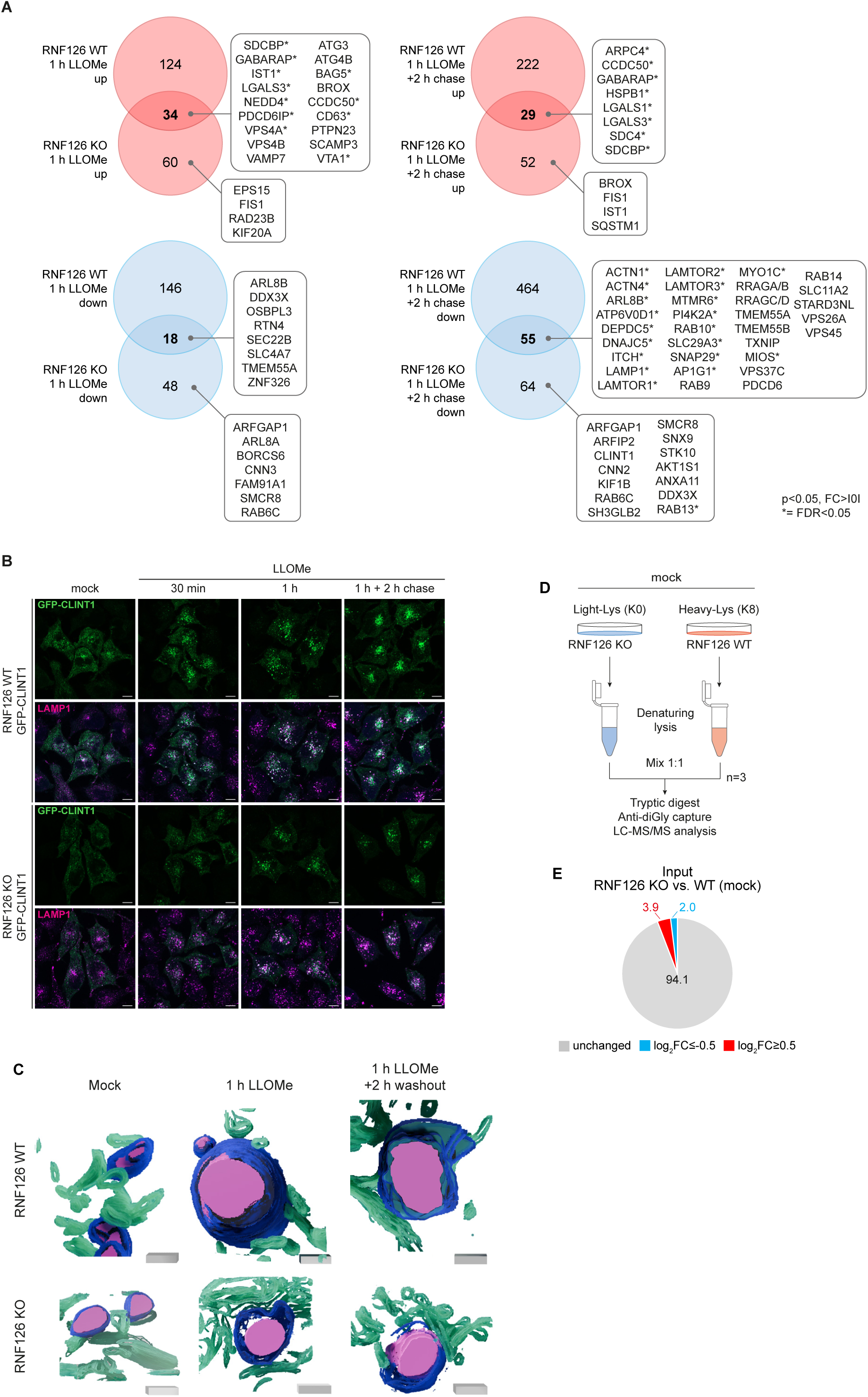
RNF126 loss affects lysosomal proteome in response to damage. (A) Venn diagram of proteins significantly enriched or decreased in the proximity of lysosomes (p-value<0.05, FC>0, *= FDR<0.05) in RNF126 WT or KO cells differentially treated with 1h LLOMe or 1 h LLOMe followed by 2 h chase. Selected common and RNF16 KO specific candidates are highlighted. (B) RNF126 WT and KO cells expressing GFP-CLINT1 were treated as indicated, fixed and stained with an anti-LAMP1 antibody. Scale bars, 10 µm. (C) Electron tomogram of RNF126 WT and KO cells treated as indicated from Figure 7G. Lysosomal dense core is colored pink, limiting lysosomal membrane blue and surrounding ER green. Scale cubes, length 200 nm. (D) Experimental workflow of SILAC-based diGly MS in RNF126 WT and KO cells. Experiments were performed in biological triplicates and statistical analysis using one sample Student’s t-test (p-value<0.05) with multiple comparison correction (Benjamini-Hochberg correction FDR<0.05). (E) Percentage of quantified proteins whose abundance changed by at least twofold (log_2_ SILAC ratio >1 or <-1).s

## REFERENCES

1. Papadopoulos, C., and Meyer, H. (2017). Detection and Clearance of Damaged Lysosomes by the Endo-Lysosomal Damage Response and Lysophagy. Current biology : CB 27, R1330–R1341. 10.1016/j.cub.2017.11.012.

2. Wang, F., Gómez-Sintes, R., and Boya, P. (2018). Lysosomal membrane permeabilization and cell death. Traffic (Copenhagen, Denmark) 19, 918–931. 10.1111/tra.12613.

3. Gómez-Sintes, R., Ledesma, M.D., and Boya, P. (2016). Lysosomal cell death mechanisms in aging. Ageing research reviews 32, 150–168. 10.1016/j.arr.2016.02.009.

4. Zoncu, R., and Perera, R.M. (2022). Built to last: lysosome remodeling and repair in health and disease. Trends in cell biology 32, 597–610. 10.1016/j.tcb.2021.12.009.

5. Yang, H., and Tan, J.X. (2023). Lysosomal quality control: molecular mechanisms and therapeutic implications. Trends in cell biology 33, 749–764. 10.1016/j.tcb.2023.01.001.

6. Meyer, H., and Kravic, B. (2024). The Endo-Lysosomal Damage Response. Annual review of biochemistry 93, 367–387. 10.1146/annurev-biochem-030222-102505.

7. Skowyra, M.L., Schlesinger, P.H., Naismith, T.V., and Hanson, P.I. (2018). Triggered recruitment of ESCRT machinery promotes endolysosomal repair. Science (New York, N.Y.) 360. 10.1126/science.aar5078.

8. Radulovic, M., Schink, K.O., Wenzel, E.M., Nähse, V., Bongiovanni, A., Lafont, F., and Stenmark, H. (2018). ESCRT-mediated lysosome repair precedes lysophagy and promotes cell survival. The EMBO journal 37. 10.15252/embj.201899753.

9. Jia, J., Claude-Taupin, A., Gu, Y., Choi, S.W., Peters, R., Bissa, B., Mudd, M.H., Allers, L., Pallikkuth, S., and Lidke, K.A., et al. (2020). Galectin-3 Coordinates a Cellular System for Lysosomal Repair and Removal. Developmental cell 52, 69–87.e8. 10.1016/j.devcel.2019.10.025.

10. Herbst, S., Campbell, P., Harvey, J., Bernard, E.M., Papayannopoulos, V., Wood, N.W., Morris, H.R., and Gutierrez, M.G. (2020). LRRK2 activation controls the repair of damaged endomembranes in macrophages. The EMBO journal 39, e104494. 10.15252/embj.2020104494.

11. Radulovic, M., Wenzel, E.M., Gilani, S., Holland, L.K., Lystad, A.H., Phuyal, S., Olkkonen, V.M., Brech, A., Jäättelä, M., and Maeda, K., et al. (2022). Cholesterol transfer via endoplasmic reticulum contacts mediates lysosome damage repair. The EMBO journal 41, e112677. 10.15252/embj.2022112677.

12. Tan, J.X., and Finkel, T. (2022). A phosphoinositide signalling pathway mediates rapid lysosomal repair. Nature 609, 815–821. 10.1038/s41586-022-05164-4.

13. Jia, J., Abudu, Y.P., Claude-Taupin, A., Gu, Y., Kumar, S., Choi, S.W., Peters, R., Mudd, M.H., Allers, L., and Salemi, M., et al. (2018). Galectins Control mTOR in Response to Endomembrane Damage. Molecular cell 70, 120–135.e8. 10.1016/j.molcel.2018.03.009.

14. Nakamura, S., Shigeyama, S., Minami, S., Shima, T., Akayama, S., Matsuda, T., Esposito, A., Napolitano, G., Kuma, A., and Namba-Hamano, T., et al. (2020). LC3 lipidation is essential for TFEB activation during the lysosomal damage response to kidney injury. Nature cell biology 22, 1252–1263. 10.1038/s41556-020-00583-9.

15. Ikuko Maejima, Atsushi Takahashi, Hiroko Omori, Tomonori Kimura, Yoshitsugu Takabatake, Tatsuya Saitoh, Akitsugu Yamamoto, Maho Hamasaki, Takeshi Noda, and Yoshitaka Isaka, et al. Autophagy sequesters damaged lysosomes to control lysosomal biogenesis and kidney injury.

16. Hung, Y.-H., Chen, L.M.-W., Yang, J.-Y., and Yang, W.Y. (2013). Spatiotemporally controlled induction of autophagy-mediated lysosome turnover. Nature communications 4, 2111. 10.1038/ncomms3111.

17. Hoyer, M.J., Swarup, S., and Harper, J.W. (2022). Mechanisms Controlling Selective Elimination of Damaged Lysosomes. Current opinion in physiology 29. 10.1016/j.cophys.2022.100590.

18. Koerver, L., Papadopoulos, C., Liu, B., Kravic, B., Rota, G., Brecht, L., Veenendaal, T., Polajnar, M., Bluemke, A., and Ehrmann, M., et al. (2019). The ubiquitin-conjugating enzyme UBE2QL1 coordinates lysophagy in response to endolysosomal damage. EMBO reports 20, e48014. 10.15252/embr.201948014.

19. Yoshida, Y., Yasuda, S., Fujita, T., Hamasaki, M., Murakami, A., Kawawaki, J., Iwai, K., Saeki, Y., Yoshimori, T., and Matsuda, N., et al. (2017). Ubiquitination of exposed glycoproteins by SCFFBXO27 directs damaged lysosomes for autophagy. Proceedings of the National Academy of Sciences of the United States of America 114, 8574–8579. 10.1073/pnas.1702615114.

20. Teranishi, H., Tabata, K., Saeki, M., Umemoto, T., Hatta, T., Otomo, T., Yamamoto, K., Natsume, T., Yoshimori, T., and Hamasaki, M. (2022). Identification of CUL4A-DDB1-WDFY1 as an E3 ubiquitin ligase complex involved in initiation of lysophagy. Cell reports 40, 111349. 10.1016/j.celrep.2022.111349.

21. Kravić, B., Bionda, T., Siebert, A., Gahlot, P., Levantovsky, S., Behrends, C., and Meyer, H. (2022). Ubiquitin profiling of lysophagy identifies actin stabilizer CNN2 as a target of VCP/p97 and uncovers a link to HSPB1. Molecular cell 82, 2633–2649.e7. 10.1016/j.molcel.2022.06.012.

22. Cross, J., Durgan, J., McEwan, D.G., Tayler, M., Ryan, K.M., and Florey, O. (2023). Lysosome damage triggers direct ATG8 conjugation and ATG2 engagement via non-canonical autophagy. The Journal of cell biology 222, e202303078. 10.1083/jcb.202303078.

23. Bonet-Ponce, L., Beilina, A., Williamson, C.D., Lindberg, E., Kluss, J.H., Saez-Atienzar, S., Landeck, N., Kumaran, R., Mamais, A., and Bleck, C.K.E., et al. (2020). LRRK2 mediates tubulation and vesicle sorting from lysosomes. Science advances 6. 10.1126/sciadv.abb2454.

24. Bhattacharya, A., Mukherjee, R., Kuncha, S.K., Brunstein, M.E., Rathore, R., Junek, S., Münch, C., and Dikic, I. (2023). A lysosome membrane regeneration pathway depends on TBC1D15 and autophagic lysosomal reformation proteins. Nature cell biology 25, 685–698. 10.1038/s41556-023-01125-9.

25. Lee, C., Lamech, L., Johns, E., and Overholtzer, M. (2020). Selective Lysosome Membrane Turnover Is Induced by Nutrient Starvation. Developmental cell 55, 289–297.e4. 10.1016/j.devcel.2020.08.008.

26. Ogura, M., Kaminishi, T., Shima, T., Torigata, M., Bekku, N., Tabata, K., Minami, S., Nishino, K., Nezu, A., and Hamasaki, M., et al. (2023). Microautophagy regulated by STK38 and GABARAPs is essential to repair lysosomes and prevent aging. EMBO reports 24, e57300. 10.15252/embr.202357300.

27. Gahlot, P., Kravic, B., Rota, G., van den Boom, J., Levantovsky, S., Schulze, N., Maspero, E., Polo, S., Behrends, C., and Meyer, H. (2024). Lysosomal damage sensing and lysophagy initiation by SPG20-ITCH. Molecular cell 84, 1556–1569.e10. 10.1016/j.molcel.2024.02.029.

28. Hung, V., Udeshi, N.D., Lam, S.S., Loh, K.H., Cox, K.J., Pedram, K., Carr, S.A., and Ting, A.Y. (2016). Spatially resolved proteomic mapping in living cells with the engineered peroxidase APEX2. Nature protocols 11, 456–475. 10.1038/nprot.2016.018.

29. Thiele, D.L., and Lipsky, P.E. (1990). Mechanism of L-leucyl-L-leucine methyl ester-mediated killing of cytotoxic lymphocytes: dependence on a lysosomal thiol protease, dipeptidyl peptidase I, that is enriched in these cells. Proceedings of the National Academy of Sciences of the United States of America 87, 83–87. 10.1073/pnas.87.1.83.

30. Uchimoto, T., Nohara, H., Kamehara, R., Iwamura, M., Watanabe, N., and Kobayashi, Y. (1999). Mechanism of apoptosis induced by a lysosomotropic agent, L-Leucyl-L-Leucine methyl ester. Apoptosis : an international journal on programmed cell death 4, 357–362. 10.1023/a:1009695221038.

31. Papadopoulos, C., Kirchner, P., Bug, M., Grum, D., Koerver, L., Schulze, N., Poehler, R., Dressler, A., Fengler, S., and Arhzaouy, K., et al. (2017). VCP/p97 cooperates with YOD1, UBXD1 and PLAA to drive clearance of ruptured lysosomes by autophagy. The EMBO journal 36, 135–150. 10.15252/embj.201695148.

32. Eapen, V.V., Swarup, S., Hoyer, M.J., Paulo, J.A., and Harper, J.W. (2021). Quantitative proteomics reveals the selectivity of ubiquitin-binding autophagy receptors in the turnover of damaged lysosomes by lysophagy. eLife 10. 10.7554/eLife.72328.

33. Chauhan, S., Kumar, S., Jain, A., Ponpuak, M., Mudd, M.H., Kimura, T., Choi, S.W., Peters, R., Mandell, M., and Bruun, J.-A., et al. (2016). TRIMs and Galectins Globally Cooperate and TRIM16 and Galectin-3 Co-direct Autophagy in Endomembrane Damage Homeostasis. Developmental cell 39, 13–27. 10.1016/j.devcel.2016.08.003.

34. Zhang, W., Yang, X., Chen, L., Liu, Y.-Y., Venkatarangan, V., Reist, L., Hanson, P., Xu, H., Wang, Y., and Li, M. (2021). A conserved ubiquitin-and ESCRT-dependent pathway internalizes human lysosomal membrane proteins for degradation. PLoS biology 19, e3001361. 10.1371/journal.pbio.3001361.

35. Abu-Remaileh, M., Wyant, G.A., Kim, C., Laqtom, N.N., Abbasi, M., Chan, S.H., Freinkman, E., and Sabatini, D.M. (2017). Lysosomal metabolomics reveals V-ATPase-and mTOR-dependent regulation of amino acid efflux from lysosomes. Science (New York, N.Y.) 358, 807–813. 10.1126/science.aan6298.

36. Repnik, U., Borg Distefano, M., Speth, M.T., Ng, M.Y.W., Progida, C., Hoflack, B., Gruenberg, J., and Griffiths, G. (2017). L-leucyl-L-leucine methyl ester does not release cysteine cathepsins to the cytosol but inactivates them in transiently permeabilized lysosomes. Journal of cell science 130, 3124–3140. 10.1242/jcs.204529.

37. Berg, T.O., Strømhaug, E., Løvdal, T., Seglen, O., and Berg, T. (1994). Use of glycyl-L-phenylalanine 2-naphthylamide, a lysosome-disrupting cathepsin C substrate, to distinguish between lysosomes and prelysosomal endocytic vacuoles. Biochemical Journal 300 ( *Pt 1*), 229–236. 10.1042/bj3000229.

38. Florey, O., Gammoh, N., Kim, S.E., Jiang, X., and Overholtzer, M. (2015). V-ATPase and osmotic imbalances activate endolysosomal LC3 lipidation. Autophagy 11, 88–99. 10.4161/15548627.2014.984277.

39. Jacquin, E., Leclerc-Mercier, S., Judon, C., Blanchard, E., Fraitag, S., and Florey, O. (2017). Pharmacological modulators of autophagy activate a parallel noncanonical pathway driving unconventional LC3 lipidation. Autophagy 13, 854–867. 10.1080/15548627.2017.1287653.

40. Durgan, J., Lystad, A.H., Sloan, K., Carlsson, S.R., Wilson, M.I., Marcassa, E., Ulferts, R., Webster, J., Lopez-Clavijo, A.F., and Wakelam, M.J., et al. (2021). Non-canonical autophagy drives alternative ATG8 conjugation to phosphatidylserine. Molecular cell 81, 2031–2040.e8. 10.1016/j.molcel.2021.03.020.

41. van der Paal, J., Neyts, E.C., Verlackt, C.C.W., and Bogaerts, A. (2016). Effect of lipid peroxidation on membrane permeability of cancer and normal cells subjected to oxidative stress. Chemical science 7, 489–498. 10.1039/c5sc02311d.

42. Bour, A., Kruglik, S.G., Chabanon, M., Rangamani, P., Puff, N., and Bonneau, S. (2019). Lipid Unsaturation Properties Govern the Sensitivity of Membranes to Photoinduced Oxidative Stress. Biophysical journal 116, 910–920. 10.1016/j.bpj.2019.01.033.

43. Bhardwaj, M., Lee, J.J., Versace, A.M., Harper, S.L., Goldman, A.R., Crissey, M.A.S., Jain, V., Singh, M.P., Vernon, M., and Aplin, A.E., et al. (2023). Lysosomal lipid peroxidation regulates tumor immunity. The Journal of clinical investigation 133. 10.1172/JCI164596.

44. Shi, X., Mao, Y., Daniel, L.N., Saffiotti, U., Dalal, N.S., and Vallyathan, V. (1994). Silica radical-induced DNA damage and lipid peroxidation. Environmental health perspectives 102 *Suppl 10*, 149–154. 10.1289/ehp.94102s10149.

45. Zhi, X., Zhao, D., Wang, Z., Zhou, Z., Wang, C., Chen, W., Liu, R., and Chen, C. (2013). E3 ubiquitin ligase RNF126 promotes cancer cell proliferation by targeting the tumor suppressor p21 for ubiquitin-mediated degradation. Cancer research 73, 385–394. 10.1158/0008-5472.CAN-12-0562.

46. Liu, W., Zheng, M., Zhang, R., Jiang, Q., Du, G., Wu, Y., Yang, C., Li, F., Li, W., and Wang, L., et al. (2023). RNF126-Mediated MRE11 Ubiquitination Activates the DNA Damage Response and Confers Resistance of Triple-Negative Breast Cancer to Radiotherapy. Advanced Science 10, e2203884. 10.1002/advs.202203884.

47. Rodrigo-Brenni, M.C., Gutierrez, E., and Hegde, R.S. (2014). Cytosolic quality control of mislocalized proteins requires RNF126 recruitment to Bag6. Molecular cell 55, 227–237. 10.1016/j.molcel.2014.05.025.

48. Hu, X., Wang, L., Wang, Y., Ji, J., Li, J., Wang, Z., Li, C., Zhang, Y., and Zhang, Z.-R. (2020). RNF126-Mediated Reubiquitination Is Required for Proteasomal Degradation of p97-Extracted Membrane Proteins. Molecular cell 79, 320–331.e9. 10.1016/j.molcel.2020.06.023.

49. Krysztofinska, E.M., Martínez-Lumbreras, S., Thapaliya, A., Evans, N.J., High, S., and Isaacson, R.L. (2016). Structural and functional insights into the E3 ligase, RNF126. Scientific reports 6, 26433. 10.1038/srep26433.

50. Jia, J., Wang, F., Bhujabal, Z., Peters, R., Mudd, M., Duque, T., Allers, L., Javed, R., Salemi, M., and Behrends, C., et al. (2022). Stress granules and mTOR are regulated by membrane atg8ylation during lysosomal damage. The Journal of cell biology 221. 10.1083/jcb.202207091.

51. Bussi, C., Mangiarotti, A., Vanhille-Campos, C., Aylan, B., Pellegrino, E., Athanasiadi, N., Fearns, A., Rodgers, A., Franzmann, T.M., and Šarić, A., et al. (2023). Stress granules plug and stabilize damaged endolysosomal membranes. Nature 623, 1062–1069. 10.1038/s41586-023-06726-w.

52. Yasar Arfat T. Kasu, Akshaya Arva, Jess Johnson, Christin Sajan, Jasmin Manzano, Andrew Hennes, Jacy Haynes, and Christopher S. Brower. BAG6 prevents the aggregation of neurodegeneration-associated fragments of TDP43.

53. Jia, P., Tian, T., Li, Z., Wang, Y., Lin, Y., Zeng, W., Ye, Y., He, M., Ni, X., and Pan, J.’a., et al. (2023). CCDC50 promotes tumor growth through regulation of lysosome homeostasis. EMBO reports 24, e56948. 10.15252/embr.202356948.

54. Milkereit, R., and Rotin, D. (2011). A role for the ubiquitin ligase Nedd4 in membrane sorting of LAPTM4 proteins. PloS one 6, e27478. 10.1371/journal.pone.0027478.

55. Smith, C.J., Berry, D.M., and McGlade, C.J. (2013). The E3 ubiquitin ligases RNF126 and Rabring7 regulate endosomal sorting of the epidermal growth factor receptor. Journal of cell science 126, 1366–1380. 10.1242/jcs.116129.

56. Smith, C.J., and McGlade, C.J. (2014). The ubiquitin ligase RNF126 regulates the retrograde sorting of the cation-independent mannose 6-phosphate receptor. Experimental cell research 320, 219–232. 10.1016/j.yexcr.2013.11.013.

57. Milkereit, R., Persaud, A., Vanoaica, L., Guetg, A., Verrey, F., and Rotin, D. (2015). LAPTM4b recruits the LAT1-4F2hc Leu transporter to lysosomes and promotes mTORC1 activation. Nature communications 6, 7250. 10.1038/ncomms8250.

58. Jia, Y., Jia, R., Dai, Z., Zhou, J., Ruan, J., Chng, W., Cai, Z., and Zhang, X. (2024). Stress granules in cancer: Adaptive dynamics and therapeutic implications. iScience 27, 110359. 10.1016/j.isci.2024.110359.

59. Saint-Pol, A., Yélamos, B., Amessou, M., Mills, I.G., Dugast, M., Tenza, D., Schu, P., Antony, C., McMahon, H.T., and Lamaze, C., et al. (2004). Clathrin adaptor epsinR is required for retrograde sorting on early endosomal membranes. Developmental cell 6, 525–538. 10.1016/s1534-5807(04)00100-5.

60. Mills, I.G., Praefcke, G.J.K., Vallis, Y., Peter, B.J., Olesen, L.E., Gallop, J.L., Butler, P.J.G., Evans, P.R., and McMahon, H.T. (2003). EpsinR: an AP1/clathrin interacting protein involved in vesicle trafficking. The Journal of cell biology 160, 213–222. 10.1083/jcb.200208023.

61. Chidambaram, S., Müllers, N., Wiederhold, K., Haucke, V., and Mollard, G.F. von (2004). Specific interaction between SNAREs and epsin N-terminal homology (ENTH) domains of epsin-related proteins in trans-Golgi network to endosome transport. The Journal of biological chemistry 279, 4175–4179. 10.1074/jbc.M308667200.

62. Miller, S.E., Collins, B.M., McCoy, A.J., Robinson, M.S., and Owen, D.J. (2007). A SNARE-adaptor interaction is a new mode of cargo recognition in clathrin-coated vesicles. Nature 450, 570–574. 10.1038/nature06353.

63. Nachef, M., Ali, A.K., Almutairi, S.M., and Lee, S.-H. (2021). Targeting SLC1A5 and SLC3A2/SLC7A5 as a Potential Strategy to Strengthen Anti-Tumor Immunity in the Tumor Microenvironment. Frontiers in Immunology 12, 624324. 10.3389/fimmu.2021.624324.

64. Zellner, S., Schifferer, M., and Behrends, C. (2021). Systematically defining selective autophagy receptor-specific cargo using autophagosome content profiling. Molecular cell 81, 1337–1354.e8. 10.1016/j.molcel.2021.01.009.

65. Bocchini, V., Mazzolla, R., Barluzzi, R., Blasi, E., Sick, P., and Kettenmann, H. (1992). An immortalized cell line expresses properties of activated microglial cells. Journal of neuroscience research 31, 616–621. 10.1002/jnr.490310405.

66. Ran, F.A., Hsu, P.D., Wright, J., Agarwala, V., Scott, D.A., and Zhang, F. (2013). Genome engineering using the CRISPR-Cas9 system. Nat Protoc 8, 2281–2308. 10.1038/nprot.2013.143.

67. Wefers, B., Wurst, W., and Kühn, R. (2023). Gene Editing in Mouse Zygotes Using the CRISPR/Cas9 System. Methods in molecular biology (Clifton, N.J.) 2631, 207–230. 10.1007/978-1-0716-2990-1_8.

68. Concordet, J.-P., and Haeussler, M. (2018). CRISPOR: intuitive guide selection for CRISPR/Cas9 genome editing experiments and screens. Nucleic acids research 46, W242–W245. 10.1093/nar/gky354.

69. Sograte-Idrissi, S., Oleksiievets, N., Isbaner, S., Eggert-Martinez, M., Enderlein, J., Tsukanov, R., and Opazo, F. (2019). Nanobody Detection of Standard Fluorescent Proteins Enables Multi-Target DNA-PAINT with High Resolution and Minimal Displacement Errors. Cells 8. 10.3390/cells8010048.

70. Jungmann, R., Avendaño, M.S., Woehrstein, J.B., Dai, M., Shih, W.M., and Yin, P. (2014). Multiplexed 3D cellular super-resolution imaging with DNA-PAINT and Exchange-PAINT. Nature methods 11, 313–318. 10.1038/nmeth.2835.

71. Schnitzbauer, J., Strauss, M.T., Schlichthaerle, T., Schueder, F., and Jungmann, R. (2017). Super-resolution microscopy with DNA-PAINT. Nature protocols 12, 1198–1228. 10.1038/nprot.2017.024.

72. Mastronarde, D.N. (2005). Automated electron microscope tomography using robust prediction of specimen movements. Journal of structural biology 152, 36–51. 10.1016/j.jsb.2005.07.007.

73. Kremer, J.R., Mastronarde, D.N., and McIntosh, J.R. (1996). Computer visualization of three-dimensional image data using IMOD. Journal of structural biology 116, 71–76. 10.1006/jsbi.1996.0013.

74. Berger, D.R., Seung, H.S., and Lichtman, J.W. (2018). VAST (Volume Annotation and Segmentation Tool): Efficient Manual and Semi-Automatic Labeling of Large 3D Image Stacks. Frontiers in neural circuits 12, 88. 10.3389/fncir.2018.00088.

75. V. Eapen, V., Swarup, S., Hoyer, M., and JW, H. (2021). Analysis of Lysophagic Flux in Cultured Cells using Lyso-Keima v2.

76. Tanaka, A., Cleland, M.M., Xu, S., Narendra, D.P., Suen, D.-F., Karbowski, M., and Youle, R.J. (2010). Proteasome and p97 mediate mitophagy and degradation of mitofusins induced by Parkin. The Journal of cell biology 191, 1367–1380. 10.1083/jcb.201007013.

77. Lobingier, B.T., Hüttenhain, R., Eichel, K., Miller, K.B., Ting, A.Y., Zastrow, M. von, and Krogan, N.J. (2017). An Approach to Spatiotemporally Resolve Protein Interaction Networks in Living Cells. Cell 169, 350–360.e12. 10.1016/j.cell.2017.03.022.

78. Rappsilber, J., Ishihama, Y., and Mann, M. (2003). Stop and go extraction tips for matrix-assisted laser desorption/ionization, nanoelectrospray, and LC/MS sample pretreatment in proteomics. Analytical chemistry 75, 663–670. 10.1021/ac026117i.

79. Fiskin, E., Bionda, T., Dikic, I., and Behrends, C. (2016). Global Analysis of Host and Bacterial Ubiquitinome in Response to Salmonella Typhimurium Infection. Molecular cell 62, 967–981. 10.1016/j.molcel.2016.04.015.

80. Villén, J., and Gygi, S.P. (2008). The SCX/IMAC enrichment approach for global phosphorylation analysis by mass spectrometry. Nature protocols 3, 1630–1638. 10.1038/nprot.2008.150.

81. Cox, J., and Mann, M. (2008). MaxQuant enables high peptide identification rates, individualized p.p.b.-range mass accuracies and proteome-wide protein quantification. Nature biotechnology 26, 1367–1372. 10.1038/nbt.1511.

82. Cox, J., Neuhauser, N., Michalski, A., Scheltema, R.A., Olsen, J.V., and Mann, M. (2011). Andromeda: a peptide search engine integrated into the MaxQuant environment. Journal of proteome research 10, 1794–1805. 10.1021/pr101065j.

83. Tyanova, S., Temu, T., Sinitcyn, P., Carlson, A., Hein, M.Y., Geiger, T., Mann, M., and Cox, J. (2016). The Perseus computational platform for comprehensive analysis of (prote)omics data. Nature methods 13, 731–740. 10.1038/nmeth.3901.

84. Huang, D.W., Sherman, B.T., Tan, Q., Kir, J., Liu, D., Bryant, D., Guo, Y., Stephens, R., Baseler, M.W., and Lane, H.C., et al. (2007). DAVID Bioinformatics Resources: expanded annotation database and novel algorithms to better extract biology from large gene lists. Nucleic acids research 35, W169–75. 10.1093/nar/gkm415.

85. Huang, D.W., Sherman, B.T., and Lempicki, R.A. (2009). Systematic and integrative analysis of large gene lists using DAVID bioinformatics resources. Nat Protoc 4, 44–57. 10.1038/nprot.2008.211.

